# CDX2 Drives Divergent Regional Programs During Intestinal Development and Redeploys Them During Regeneration

**DOI:** 10.64898/2026.09.29.755502

**Authors:** A Lorzadeh, G Ye, M Romero-Wolf, ML Quiloan, S Sharma, E Sanchez, U Jadhav

**Author notes:** **Corresponding author:** Unmesh Jadhav, PhD, Department of Stem Cell Biology and Regenerative Medicine, Keck School of Medicine, University of Southern California, Broad CIRM Center, 1425 San Pablo Street BCC-511, Los Angeles, CA 90033. These authors contributed equally to this work (A Lorzadeh, G Ye).

## Abstract

Region-specific transcription factors (TFs) establish small intestinal and colonic identity, yet what activates their region-specific expression during development is unresolved. Moreover, discrete fetal gene sets reactivate during post-injury regeneration, but their full scope, and what reactivates the programs required to restore regional identity and tissue structure, remain unclear. By determining CDX2 and partner TF binding, enhancer activity, and gene expression dynamics across intestinal regions, in stem and differentiating cells, and through development and adult regeneration, here we show that CDX2, despite uniform expression across the fetal and adult intestinal epithelium, organizes a hierarchical enhancer network. This network cooperates with partner factors such as CTCF to establish a shared, plastic fetal enhancer landscape, which resolves into region-specific programs upstream of regional TFs. During regeneration, rather than reactivating a general fetal state, CDX2 selectively rebinds many region-specific and common enhancers for genes associated with fetal cell-motility and tissue-morphogenesis signatures along with a more recently active adult stem-cell program, each deployed across distinct crypt tiers. CDX2 thus functions as a central, context-responsive regulator, deploying fetal, adult stem-cell differentiation, and regenerative programs as tissue context requires.

**Graphical Abstract:** 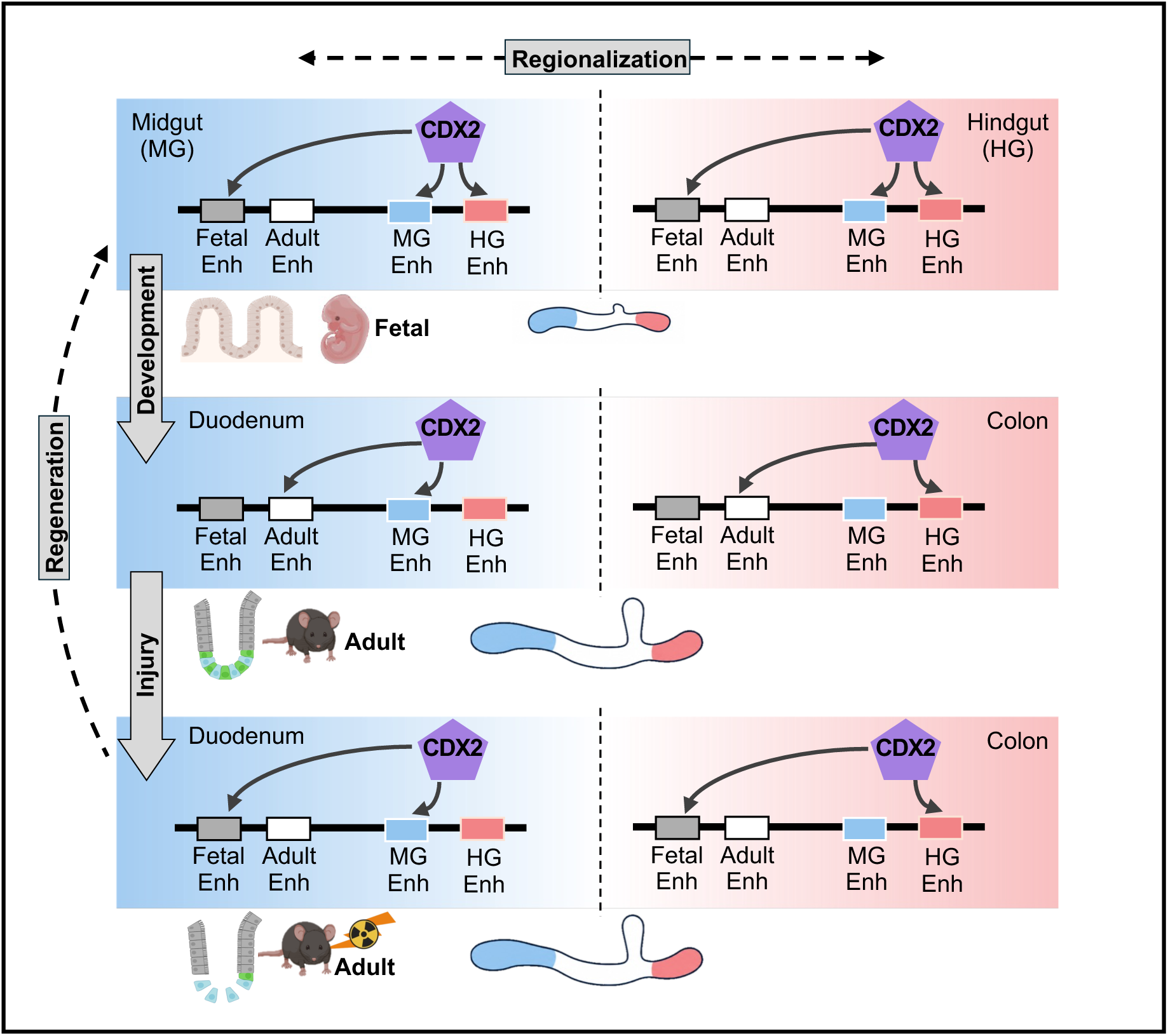

## INTRODUCTION

The formation of complex organs from common embryonic progenitors requires remarkable spatial precision, generating tissues with distinct architecture and function from a shared point of origin. In the mammalian intestine, this process regionalizes an initially plastic fetal epithelium into the duodenum, ileum, and colon, each with specialized cell types, tissue architecture, and physiology^1^. These regional identities, once established, must be preserved throughout life despite continuous epithelial renewal, and faithfully rebuilt following injury — yet how regional identity first arises from a common progenitor, and how this developmental information is later accessed to guide regeneration, remain central unresolved questions in tissue biology^1,2^.

Region-specific transcription factors mark this process: GATA4 specifies proximal small-intestinal identity, SATB2 defines colonic identity, and PDX1 and other regional regulators establish additional intestinal domains^1,3–9^. Yet these factors are effectors of regional identity, not its origin — what activates their own region-specific expression in fetal progenitor cells has remained incompletely understood.

CDX2 is uniquely positioned to resolve this question. CDX2 is ubiquitously expressed throughout the developing intestinal epithelium and indispensable for intestinal specification and differentiation^10–15^, yet this broad expression poses a paradox: how can a transcription factor expressed uniformly along the intestine generate distinct regional identities? While region-restricted co-factors and repressive chromatin modifications may also contribute, enhancers offer a particularly tractable route: enhancer activity provides a combined readout of transcription factor occupancy and chromatin state, capturing how positional information is translated into divergent regulatory output by a single, broadly expressed factor ^16,17^. However, whether and how CDX2 organizes such region-specific enhancer networks has not been defined.

This developmental logic carries direct significance for regeneration. Restoring intestinal function after injury requires rebuilding not only lost cells, but the correct regional stem-cell populations, tissue architecture, and physiology established during development. Recent work shows that discrete fetal gene sets reactivate during regeneration, suggesting developmental mechanisms are redeployed after injury^2^, but whether this reflects broad reversion to a fetal state or the selective reactivation of specific developmental processes, and what molecular machinery governs this reactivation, remain unresolved.

Here, we define the CDX2-directed enhancer hierarchy that, starting from uniform binding across intestinal regions during fetal development, resolves to establish intestinal regional identity and show how this same regulatory architecture is selectively redeployed during regeneration. Rather than reverting to a broad fetal state, regenerating cells reactivate a defined set of developmentally encoded fetal and stem-cell enhancer programs with regional and spatial precision, revealing CDX2 as a context-responsive regulator that links fetal patterning to faithful adult tissue regeneration.

## RESULTS

### A largely shared fetal enhancer landscape resolves progressively into region-specific intestinal identity

During mammalian intestinal development, region-specific structure and function emerge as fetal precursors progressively resolve into committed regional identities. The hierarchical molecular decisions driving this regionalization remain incompletely defined. To define the gene-expression patterns and cellular processes underlying the intestinal epithelium’s similar fetal origins and its resolution into distinct regional derivatives in the small intestine and colon, we compared normalized gene expression (mRNA-seq) from fetal midgut (MG) at embryonic day 11.5 (E11.5), E12.5, E14.5, and E16.5 and adult differentiated duodenal villus epithelium, with the same fetal timepoints in the hindgut (HG) and adult colonic epithelium (Figure 1A, S1A) ^18,19^. Using 15,384 genes differentially expressed across any of these conditions (*q* < 0.05, 2-fold change), we conducted K-means clustering and identified 6 groups of genes with distinct expression patterns. We identified 378 genes (Group 1) with high expression in midgut that reduced in small intestine, while 1,776 (Group 3) genes were highly expressed in hindgut, reducing expression in adult colonic epithelium, revealing differences across early fetal regional programs. These included genes known to be expressed in midgut and small intestinal lineage like *Nr2e3, Ros1, Hyal5*^20–22^, and hindgut and colon expressed genes *Ier3 and Ccl3*^23,24^. A large number of genes (6,380, Group 5) showed regionally similar fetal epithelium specific gene expression, and included known cross-regional general drivers of fetal epithelial development like *Meis1*, *Id2,* and *Gpc6*^25,26^. Conversely, the small-intestinal (duodenal) and colonic epithelia gained expression of 1,184 (Group 2) and 1,474 (Group 4) genes, respectively, during development as they matured into structurally and functionally distinct adult tissues; adult tissue development was also accompanied by significant upregulation of 2,588 genes common to midgut and hindgut (Group 6). The well-known small-intestinal identity determinant *Gata4* was part of Group 2 alongside *Sis*, *Alpi*, and *Mttp*^27–29^, and the colonic identity determinant *Satb2* was part of Group 4, alongside *Noxa1*, *Lpo*, and *Ildr1*^30–32^. While known common regulators of intestinal lineages and function such as *Ascl2*, *Hnf4a*, *Car4,* and *Vdr*^1,33,34^ that play part in development as well as adult tissue function were part of Group 6. Concordant with these observations, gene ontology analysis showed that genes in adult epithelium related Groups 2, 4, and 6 were linked to absorption, secretion, digestion, and metabolism, while development related clusters (Group 1, 3, and 5) showed association with cell proliferation and differentiation, extracellular matrix reorganization, and cell-cell adhesion, reflecting the significant growth and tissue remodeling during fetal development (Figure S1B). Lastly, 1,605 differentially expressed genes showed no specific patterns among these cell states. Regional diversity thus emerges progressively: divergence begins with a small set of fetal genes (Groups 1, 3) and expands into larger differences in the adult epithelium (Groups 2, 4), yet this divergence rests on a predominantly shared developmental (Group 5) and adult (Group 6) gene-expression program across the small intestine and colon, indicating that, despite their distinct structure and function, the two regions depend on a largely conserved core of processes.

**Figure 1.**
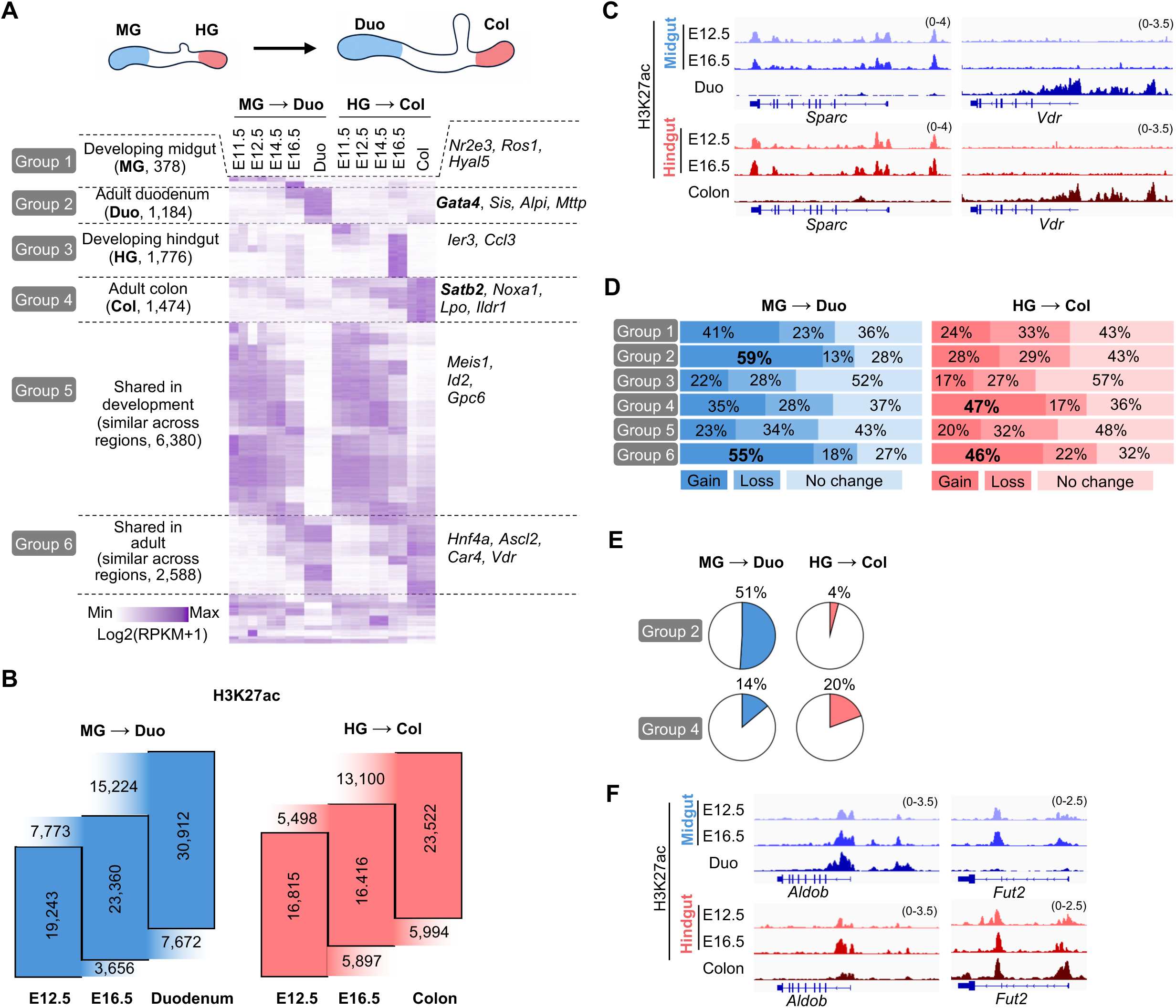
Region-specific fetal and adult identities are established through enhancer divergence. (A) Heatmap of gene-expression dynamics across fetal midgut (MG) and hindgut (HG) and their adult derivatives, duodenum and colon. Genes are grouped by expression pattern, with representative genes for each group shown alongside the heatmap. (B) Sankey diagrams illustrating transitions of fetal enhancers (E12.5 and E16.5) into adult enhancer states in the duodenum (left) and colon (right). Numbers indicate the total number of active enhancers at each developmental stage, with the number of enhancers gained and lost between consecutive stages shown above and below each transition, respectively. (C) Representative genome-browser tracks showing H3K27ac signal at fetal enhancers that are active during development (top) and adult enhancers that are acquired during maturation (bottom) along the MG→duodenum and HG→colon trajectories. (D) Quantification of enhancer fate across developmental groups, showing the proportions of enhancers that are gained, lost, or retained during MG→duodenum (blue) and HG→colon (red) differentiation. (E) Pie charts summarizing the proportion of adult small-intestinal (Group 2) and colonic (Group 4) genes associated with two or more enhancers (within ±25 kb of the transcription start site), illustrating the prevalence of multienhancer regulatory architectures. (F) Representative genome-browser tracks of H3K27ac signal at region-restricted adult genes (e.g., *Gata4*, *Satb2*), showing enhancers active in both fetal midgut and hindgut that are subsequently decommissioned in the non-corresponding region during maturation, leaving only the region-appropriate enhancer active in adult tissue.

As cis-regulatory enhancers control cell-type-specific gene expression^35,36^, we compared enhancer activation along the midgut (E12.5, E16.5)→duodenum and hindgut (E12.5, E16.5)→colon developmental trajectories. We profiled the active enhancer mark H3K27ac genome-wide by native ChIP-seq, a method suited to low cell numbers^37^, in Epcam+ epithelium and assigned enhancers to genes with a transcription start site (TSS) within 25 kb. Both regional trajectories showed extensive enhancer remodeling from fetal development into adulthood (Figure 1B, 1C). More than 11,000 fetal enhancers lost activity in adult cells of both lineages, while 21,000 and 18,000 enhancers gained significant activity (*q* < 0.05, 2X) in adult duodenum and colon, respectively, relative to their fetal counterparts. Development thus involves both gaining a large set of adult-cell-specific enhancers and systematically deactivating many fetal enhancers.

Analysis of enhancer activity gain and loss (2-fold change in H3K27ac signal) further revealed region-specific enhancer dynamics during development: 59% and 47% of enhancers linked to duodenum- and colon-specific genes (Groups 2 and 4, respectively) gained activity in adult cells, and a similar gain (55% and 46%) was seen at enhancers linked to genes commonly activated in both regions (Group 6) (Figure 1D). Conversely, 23%, 27%, and up to 34% of enhancers linked to genes highly expressed in midgut, hindgut, or shared fetal states, respectively, lost activity in adult cells, underscoring extensive fetal- and region-specific enhancer turnover (Figure 1C, D).

Beyond this temporal and regional specificity, two further features of enhancer circuits were notable. First, many genes were linked to multiple active enhancers, and enhancer number itself varied by region (Figure S1C): 51% of duodenum-specific (Group 2) genes were linked to two or more enhancers during MG→duodenum differentiation, versus only 4% during HG→colon differentiation, while 20% of colon-specific (Group 4) genes showed two or more enhancers during HG→colon differentiation, versus 14% during MG→duodenum differentiation (Figure 1E) — indicating that enhancer number itself correlates with regional expression level. Second, many genes with region-restricted activity in adult cells, including the regional determinants *Gata4* and *Satb2*, had active enhancers in both midgut and hindgut at fetal timepoints, suggesting that fetal enhancers carry an inherent cross-regional plasticity that is subsequently resolved through region-specific enhancer decommissioning during development (Figure 1F). This implies that even well-known regional determinants such as GATA4 and SATB2 require upstream, hierarchically positioned factors to direct their own temporal and region-specific activation. This reveals a nuanced logic of enhancer engagement required to establish precise, distinct regional identities across the developmental-to-adult transition: enhancers can be shared or region-specific, and can occur as single or redundant copies. This raises the question of which upstream regulators hierarchically organize this enhancer landscape as it resolves across intestinal regions during development.

### Diverging transcription factor actions build a hierarchical enhancer network, positioning CDX2 upstream of regional identity programs

Having established a hierarchical logic to enhancer engagement (Figure 1), we next asked which upstream regulators organize this hierarchy. As TF binding is a critical step in enhancer activation^16,17^, we examined the temporal dynamics of TFs linked to enhancers controlling the fetal-, adult-, and region-specific gene groups identified above. Genes with relatively higher expression in midgut and hindgut (Groups 1 and 3) were disproportionately enriched for TFs (6.6% and 6.9%, respectively) relative to the other groups, which ranged from 2.9% to 5.3% (Figure S1D), suggesting these transcription factors play an outsized role in establishing lineage-specific enhancers. TEAD and TCF factors, effectors of YAP and WNT signaling, respectively, showed higher fetal expression across both midgut and hindgut relative to their adult counterparts, consistent with their developmental role^38,39^, whereas the known regional drivers GATA4 and SATB2^1,7^ were restricted to the small intestine and colon, respectively. By contrast, HNF4A and CDX2, both with established functions across the entire intestinal tract, were expressed in both midgut and hindgut, with increasing expression in adult cells (Figure S1D). Together, these patterns suggest that diverging temporal and regional TF activity may underlie the fetal-to-adult and cross-regional differences in enhancer networks, shaping the spatiotemporal programs that establish lineage identity.

Notably, Cdx2 is expressed across all intestinal regions throughout development and adulthood, and CDX2 functions as a critical lineage-defining factor across intestinal lineages^10–14^. However, its relationship to region-specific drivers appears distinct: whereas loss of the small-intestine-specific *Gata4* or colon-specific *Satb2* disrupts cell identity within those respective regions^1,3–5,7^, deletion of *Cdx2* as early as E10 causes ectopic expression of genes characteristic of other gut-tube segments, such as the esophagus and stomach, while deletion after E12.5, or in the adult epithelium, instead disrupts maturation of functional cell types and expression of homeostatic genes^10,12,15,40,41^, together suggesting a broader, stage-dependent role for Cdx2 in regional identity that Gata4 and Satb2 do not share. While CDX2 is known to activate distinct developmental-stage-specific genes by binding distinct genomic targets across development and in adult cells^12^, whether it is directly involved in regionalization, and how it relates hierarchically to other regional drivers, has not been addressed. To test this, we integrated scRNA-seq data of 56,009 cells spanning 10 developmental timepoints, from early embryonic endoderm specification through adult intestinal epithelium from published resources and various repositories (Figure S2A)^42–44^. Cdx2 was upregulated in both midgut and hindgut endoderm beginning at E8.75, preceding the small-intestinal factors *Gata4* and *Pdx1* and the colonic factor *Satb2*, suggesting that Cdx2 may act upstream of these regional determinants (Figure 2B). This raises a key question: does *Cdx2*’s early, broad expression reflect a position upstream of regional TF networks, and if so, how is this upstream role executed at the level of individual enhancers and partner transcription factors?

**Figure 2.**
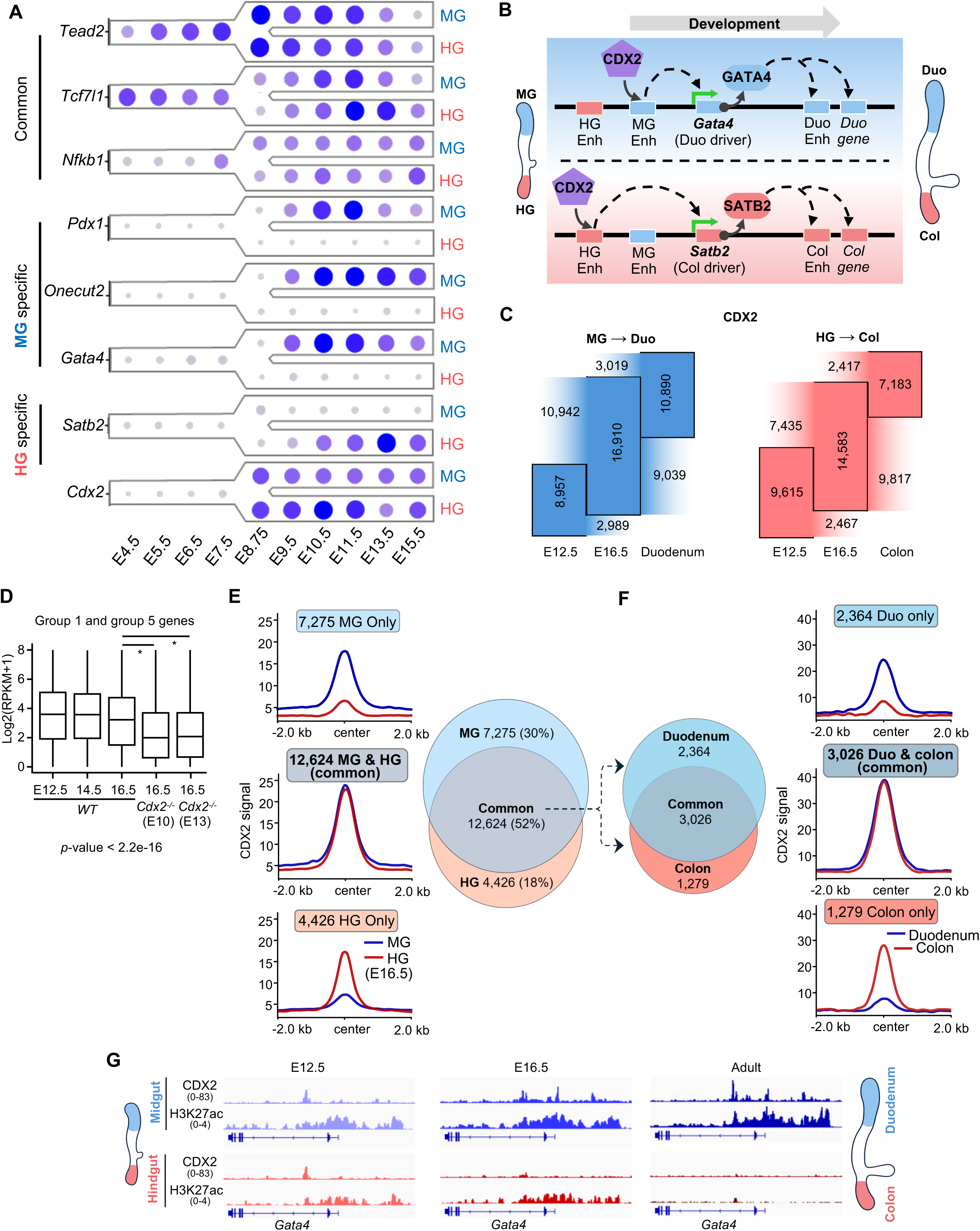
Dual roles of CDX2 binding in promoting temporal and regional intestinal identities. (A) Temporal expression dynamics of representative transcription factors during early intestinal development in the midgut (MG) and hindgut (HG). TFs are classified as common (*Tead2*, *Tcf7l1*, and *Nfkb1*), MG-specific (*Pdx1*, *Onecut2*, and *Gata4*), or HG-specific (*Satb2*). *Cdx2* is upregulated across both MG and HG lineages before the progressive emergence of region-specific transcriptional identities. (B) Schematic model of CDX2-directed regional gene activation through a hierarchical cascade along the developmental timeline: CDX2, present at both MG and HG enhancers, selectively activates the region-appropriate enhancer to drive an initial regional gene (Duo gene, top; Col gene, bottom). The resulting regional transcription factor, GATA4 (top) or SATB2 (bottom), subsequently activates additional downstream region-specific enhancers and genes, establishing regional identity. (C) Sankey diagrams depicting gain, retention, and loss of CDX2 occupancy at enhancers during MG → duodenum (blue) and HG → colon (red) development from E12.5 to adulthood. Numbers outside the boxes indicate gains and losses of CDX2 binding at the corresponding developmental transitions. (D) Expression of genes associated with enhancer Groups 1 and 5 in wild-type and CDX2-deficient intestinal epithelium. Boxplots show normalized gene expression (log2[RPKM + 1]) in wild-type epithelium at E12.5, E14.5, and E16.5, and in CDX2-deficient epithelium at E16.5 following *Cdx*2 deletion induced at either E10 or E13, demonstrating reduced expression upon *Cdx2* loss. Statistical significance was determined using a two-sided Wilcoxon signed-rank test comparing wild-type E16.5 to each *Cdx2*-deficient condition (*p* < 2.2 × 10⁻¹⁶). (E) Aggregate CDX2 occupancy profiles centered on MG-specific, shared, and HG-specifically bound sites (all unique E12.5 and E16.5 loci). The Venn diagram shows the overlap of CDX2-bound loci between MG and HG epithelium. (F) Venn diagram and aggregate CDX2 occupancy profiles showing that most fetal CDX2-bound loci shared between MG and HG resolve into distinct adult duodenal and colonic binding patterns. (G) Representative genomic locus illustrating H3K27ac and CDX2 signal at *Gata4* loci and the transition from shared fetal CDX2 occupancy to region-specific adult binding.

### Partner-TF co-occupancy and region-specific resolution mark the CDX2 binding landscape

To determine how CDX2 executes this upstream role, both in recruiting partner transcription factors and in achieving regional precision at the level of individual enhancers, we mapped genome-wide CDX2 binding in fetal (E12.5, E16.5) and adult small-intestinal and colonic epithelium. Using FACS-purified Epcam+ epithelial cells from fetal midgut and hindgut tissue and adult crypts, we performed CUT&RUN-seq and compared CDX2 binding gains and losses across hindgut → colon developmental trajectory and compared with the midgut → duodenum binding dynamics (Figure 2C)^45^. Of the 24,325 and 22,500 CDX2-binding loci identified across the small-intestinal (midgut E12.5, midgut E16.5, duodenum) and colonic (hindgut E12.5, hindgut E16.5, colon) lineages, respectively, over 78% overlapped H3K27ac+ enhancers, confirming a close relationship between CDX2 binding and enhancer activity (Figure S2B). The functional impact of this CDX2-bound enhancer activity on gene expression was further apparent in Group 1 and Group 5 genes, which include the region-specific duodenal driver *Gata4* and the developmental genes *Nkx1-2* and *Nkx2-2*. Expression of these genes was significantly reduced at E16.5 following *Cdx2* deletion at E10 or E13, relative to wild-type (Figure 2D), indicating that CDX2 binding drives enhancer activation and the downstream gene expression it controls.

Comparing CDX2 engagement across midgut and hindgut development (19,899 and 17,050 unique loci within E12.5 and E16.5, respectively), revealed that 52% of these loci overlapped between the two lineages during fetal development (Figure 2E, S2C), yet only 22% of these developmentally shared loci remained CDX2-bound in both adult duodenum and colon (Figure 2F, G, S2D). This indicates that CDX2 broadly co-binds midgut and hindgut enhancers during fetal development, followed by progressive diversification and resolution of CDX2 binding as development proceeds, ultimately yielding region-specific enhancer activity in the adult. CDX2 additionally bound loci specific to developing midgut or hindgut that were subsequently lost in adult cells, while gaining new region- and adult-specific binding elsewhere (Figure S2E), indicating that CDX2 supports both developmental-stage-specific and evolving region-specific enhancer and gene-activation programs as required. Notably, cell migration, tissue structure, and morphogenesis processes were most strongly enriched among genes commonly bound by CDX2 across both midgut and hindgut development, though also present among genes bound in either region alone (Figure S2F), suggesting that CDX2 supports a baseline structural program shared across regions, upon which region-specific developmental programs are subsequently layered.

To test whether this dynamic, stage- and region-specific binding pattern is unique to CDX2, we compared it with CTCF, a transcription factor known to bind broadly at mammalian enhancers and to play a central role in chromatin organization^17,46,47^. CTCF offers a useful contrast, as we recently showed that its recruitment is CDX2-dependent at many genomic loci while remaining CDX2-independent at others^45^. Using CTCF CUT&RUN in developing midgut and hindgut (E12.5, E16.5) and adult duodenal and colonic crypts (Figure 3A), we found that at 1,758 and 1,883 loci gaining CDX2 binding during midgut-to-duodenum and hindgut-to-colon development, respectively, CTCF binding tracked CDX2 gains, but only at loci where the two factors co-bind (Figure 3A, B). By contrast, CTCF binding at 1,539 and 2,428 CDX2-independent sites showed no consistent trend across midgut or hindgut developmental stages or their adult counterparts. Together, these results indicate that CDX2 selectively binds and activates a diverse set of enhancers and directs the developmental- and region-specific recruitment of partner factors such as CTCF.

**Figure 3.**
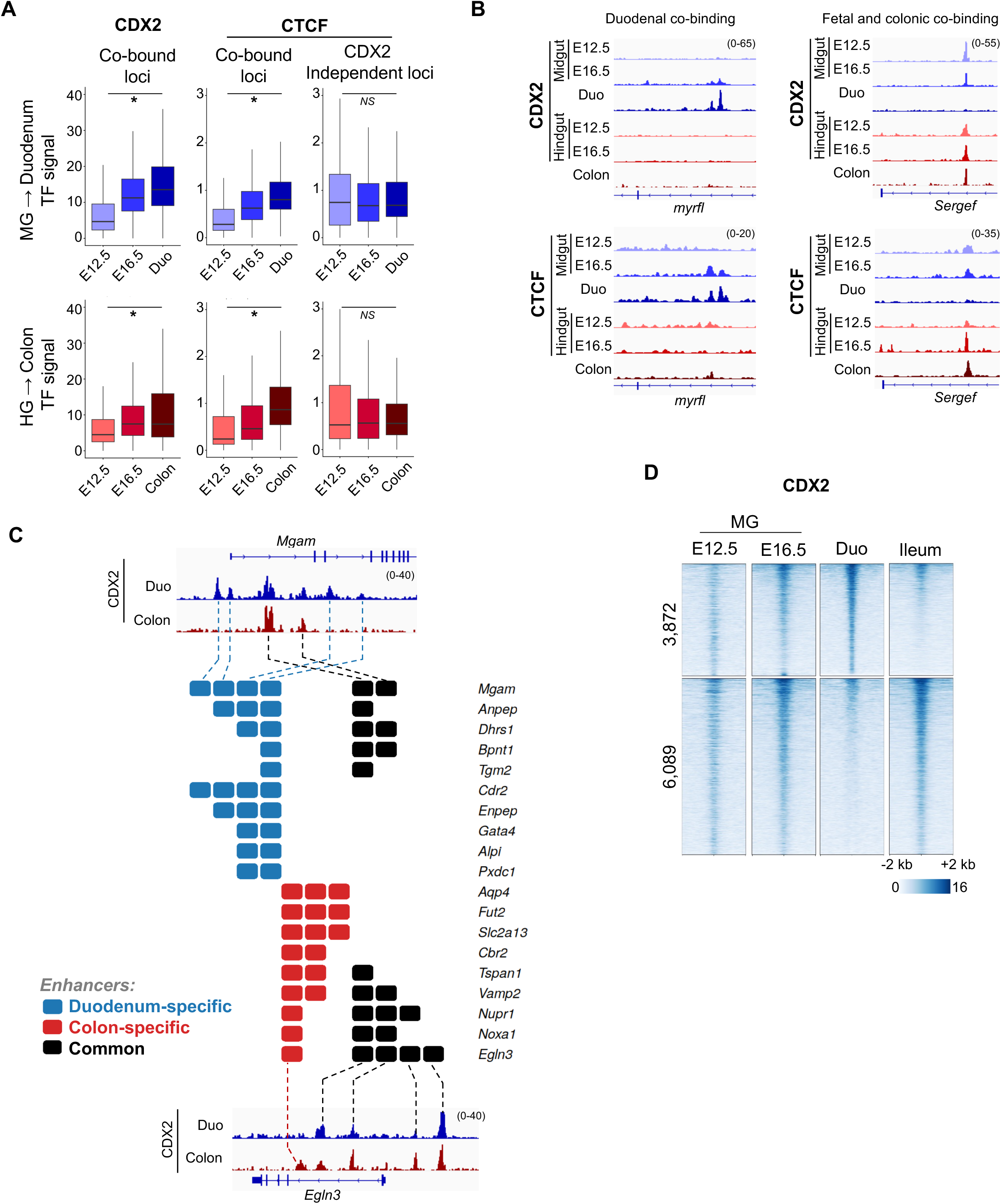
Dynamic and multienhancer CDX2 binding shapes region-specific recruitment of partner transcription factor. (A) Quantification of CDX2 occupancy and CTCF co-occupancy during intestinal development and in adult tissues, at loci gaining CDX2 binding during MG→duodenum (top; n = 1,758) and HG→colon (bottom; n = 1,883) development. Left, CDX2 signal at these loci across fetal (E12.5, E16.5) and adult (duodenum or colon) stages. Middle, CTCF signal at the same CDX2 co-bound loci. Right, CTCF signal at CDX2-independent loci (MG→duodenum, n = 1,539; HG→colon, n = 2,428), which do not gain CDX2 binding across development. Boxplots show normalized transcription-factor signal. \**p* < 2.2 × 10^-16^; NS, not significant. (B) Representative genome-browser view of CDX2 and CTCF signal showing dynamic CDX2 and CTCF occupancy across embryonic and adult intestinal tissues at a representative locus. (C) CDX2 multienhancer occupancy map for duodenal (blue) and colonic (red) genes. Each row represents one gene, and each square denotes an individual CDX2-bound enhancer located within 25 kb of the transcription start site. Black squares denote enhancers commonly bound by CDX2 in duodenum and colon. (D) Heatmaps of CDX2 occupancy showing resolution of fetal midgut binding patterns (E12.5 and E16.5) into distinct CDX2-bound loci in adult duodenum and ileum.

Notably, several region-specific genes, including the duodenal and colonic markers *Gata4* and *Satb2*, exhibited multiple and shared CDX2-bound enhancers that were active (H3K27ac+) in both regions (Figure 3C, S2D). Moreover, such multiple and shared CDX2 bound enhancers were prevalent in midgut and hindgut fetal precursors at E12.5 and E16.5 (Figure S2D, S3A). As development proceeded (E16.5 and adult), this shared binding pattern resolved: CDX2 binding was selectively retained at region-specific enhancers, while non-region-specific enhancers lost CDX2 occupancy and became inactive (H3K27ac−), and one or more adult-specific CDX2-bound enhancers were acquired, together generating distinct combinations of active, region-appropriate CDX2-bound enhancers. Overall, 1,788 genes showed at least one common enhancer in midgut and hindgut that was resolved in adult tissues. Across both developmental stage and region, a higher number of CDX2-bound enhancers correlated with higher target-gene expression (Figure 2G, S2D, S3B). Together, these data suggest that CDX2 dynamically regulates enhancer activity to direct fetal-to-adult maturation and regional identity, acting hierarchically both by activating downstream regional TF genes and by functionally shaping co-TF recruitment at appropriate target enhancers.

Given CDX2’s region-specific targeting between duodenum and colon, we asked whether CDX2 binding also distinguishes cell states within the small intestine, specifically between the proximal duodenum and distal ileum, both of which arise from midgut endoderm (Figure S3C). CUT&RUN for CDX2 and native ChIP-seq for H3K27ac in ileal epithelium, compared with duodenal data, revealed 3,872 and 6,089 loci distinctly bound by CDX2 in duodenum and ileum, respectively, all of which were CDX2-bound in the midgut (Figure 3D, S3C). This evolving binding pattern activated region-specific enhancers at known regional markers, including *Pdx1* and *Gata4* in duodenum and *Fabp6* and *Slc10a2* in ileum, further supporting that CDX2 binding evolves from a common midgut origin to establish distinct regional enhancer and gene-expression programs even within the small intestine.

Collectively, CDX2 promotes well-defined fetal, adult, and regional transcriptional programs essential for intestinal development and adult epithelial identity through precise partner-TF recruitment and highly resolved, region-specific enhancer binding (including duodenum-versus-ileum distinctions). Given recent evidence that fetal genes are reactivated during intestinal stem cell regeneration^2^, the central question this raises is whether and how CDX2 redeploys the fetal enhancer landscape it builds during development when regeneration later demands it.

### Dual reengagement of fetal and stem cell CDX2 binding signatures in non-stem crypt cells during intestinal regeneration

During adult intestinal stem cell (ISC) differentiation, TA cells retain substantial plasticity, and various TA populations can regenerate the ISC compartment following radiation- or chemically induced ISC loss^48^. This regenerative capacity has been linked to reactivation of fetal-like gene programs, including Wnt- and Yap-associated signaling^2^, yet the full breadth of these fetal programs and the molecular mechanisms governing their reactivation remain undefined.

Having established CDX2’s dynamic binding across fetal development and its molecular impact, we next asked how this fetal enhancer program resolves as cells progress into the adult ISC state and, upon homeostatic differentiation, into TA cells, and whether CDX2 binding dynamics also govern the reverse, TA-to-ISC transition that underlies regeneration following injury-induced ISC loss.

We used *Lgr5^EGFP-IRES-DTR^* mice, in which ISCs throughout the intestinal tract are EGFP-labeled^49^, and isolated crypts by standard EDTA-based isolation^50^. To capture non-stem ISC daughter cells, we depleted EGFP+ ISCs from crypt preparations by FACS^51^, then performed CDX2 CUT&RUN and mRNA-seq on the remaining cells.

To determine how CDX2 occupancy at fetal loci changes as cells mature into ISCs and TA cells, we classified genome-wide CDX2 binding across fetal (E12.5, E16.5), ISC, and TA cells by how each locus’s fetal occupancy behaved during the transition to adulthood (2-fold change): 3,620 loci lost fetal CDX2 binding (Fetal; Figure 4A), while 445 loci retained fetal CDX2 binding in adult ISCs; 1,052 loci showed CDX2 signal in TA cells similar to fetal cells (Diff 1), and 1,507 loci showed ISC- and TA-specific binding gained after the fetal period (Diff 2) (Figure 4A-B). Consistent with a functional link between CDX2 binding and target gene regulation, genes linked to these groups (TSS ± 25 kb) showed expression changes concordant with CDX2 binding dynamics (Figure 4C).

**Figure 4.**
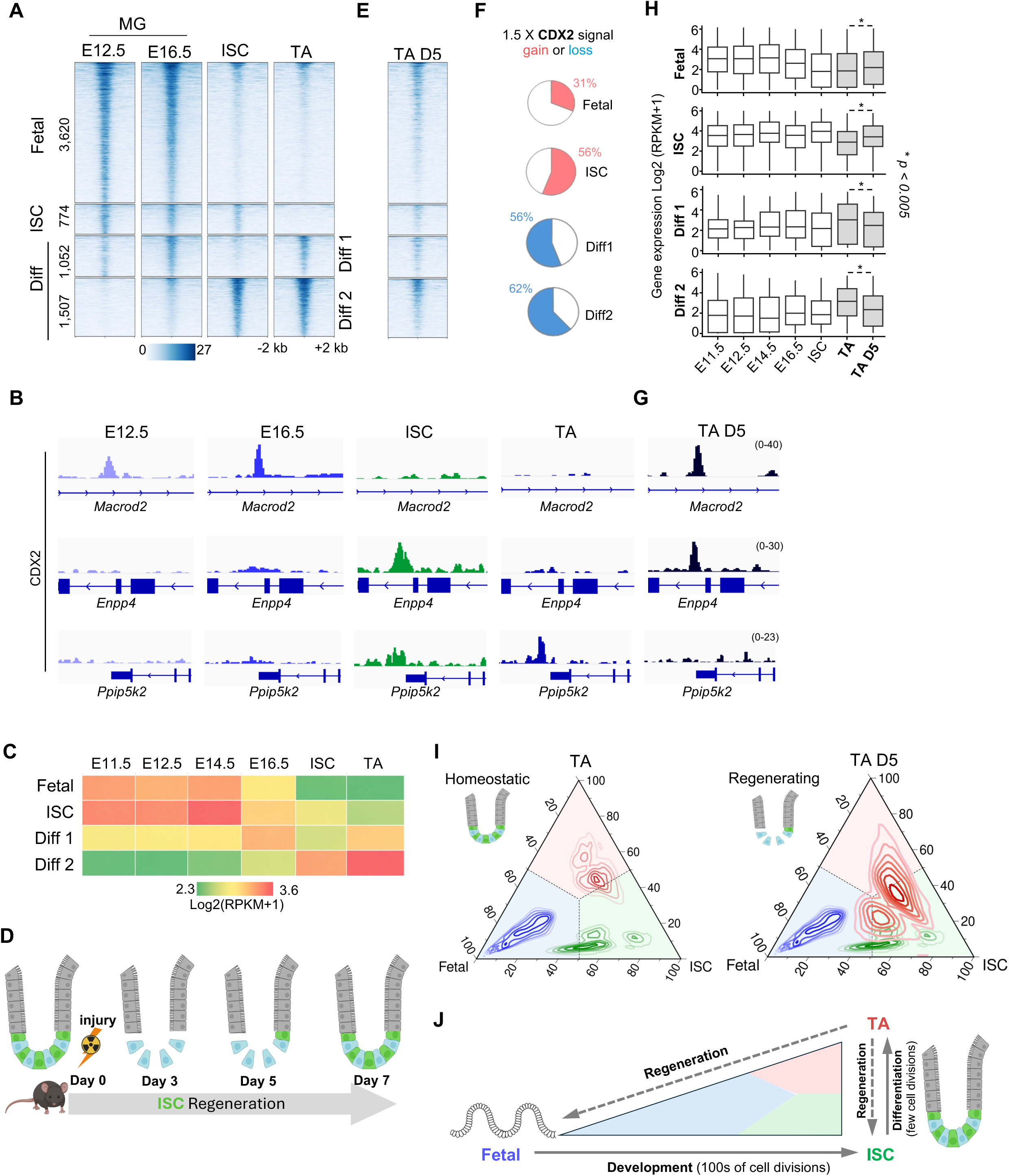
CDX2 is an on-demand transcription factor that reactivates fetal and stem-cell programs during epithelial regeneration. (A) Heatmaps showing CDX2 occupancy at fetal-, intestinal stem-cell (ISC)-, and differentiation-associated regulatory elements (Diff 1 and Diff 2) across fetal midgut (MG; E12.5 and E16.5), adult ISC, and transit-amplifying (TA) cells. CDX2 binding is dynamically redistributed among developmental and adult epithelial cell states. (B) Representative genome-browser tracks of CDX2 signal showing CDX2 occupancy at fetal-, ISC-, and differentiation-associated regulatory regions identified in panel A. (C) Heatmap representing normalized gene expression (log2[RPKM + 1]) of genes associated with the CDX2-bound regulatory groups shown in panel A across the indicated epithelial populations. (D) Schematic of radiation-induced intestinal epithelial regeneration. Following injury, Lgr5+ ISCs are depleted by day 3, reappear between days 4 and 5, and are restored in many crypts by day 7, providing a model for examining dynamic CDX2-dependent regulatory programs during regeneration. (E) Heatmap showing increased CDX2 occupancy at fetal- and ISC-associated enhancers in regenerating TA cells at day 5 post-radiation (D5 TA) compared with homeostatic TA cells, together with reduced occupancy at differentiation-associated regulatory elements. (F) Distribution of CDX2 occupancy changes during regeneration. Pie charts show the percentages of fetal- and ISC-associated loci that gain CDX2 occupancy and differentiation-associated loci that lose CDX2 occupancy by at least 1.5-fold in regenerating D5 TA cells. (G) Representative genome-browser tracks showing redistribution of CDX2 occupancy among fetal-, ISC-, and differentiation-associated regulatory regions during epithelial regeneration. (H) Expression of genes linked to the fetal-, ISC-, and differentiation-associated CDX2-bound loci defined in panel A across embryonic intestinal development, adult epithelial populations, and regenerating epithelium. Boxplots show normalized gene expression (log2[RPKM + 1]). Statistical significance was determined using the Wilcoxon signed-rank test (*p* < 0.005). (I) Ternary plots showing the relative CDX2 binding signal at fetal-, ISC-, and TA-associated loci in homeostatic TA cells (left) and TA cells isolated at day 5 post-radiation (right). Regenerating TA cells acquire a hybrid CDX2-binding state characterized by simultaneous occupancy of fetal- and ISC-associated loci, supporting a model in which CDX2 is redeployed on demand to developmental and stem-cell regulatory elements to coordinate regenerative epithelial identity. (J) Model illustrating the on-demand redeployment of CDX2 to developmental regulatory loci occupied hundreds of cell divisions earlier during fetal development and to ISC-associated loci occupied more recently in the adult stem-cell state. The simultaneous re-engagement of these temporally distinct CDX2-binding programs coordinates regenerative epithelial identity and promotes restoration of adult tissue structure and function.

With these fetal-, ISC-, and TA-defined loci established as a reference framework, we next asked how CDX2 binding in regenerating TA cells alters following ISC loss. We induced ablation of Lgr5+ ISCs in *Lgr5^EGFP-IRES-DTR^* mice via a single 10 Gy dose of abdominal X-ray radiation (Figure 4D, S4A). Lgr5+ ISCs were fully ablated by day 3 (D3) post-radiation, began re-emerging within the D4–D5 window, and were present in many crypts by day 7. We purified TA cells from D5 epithelium, depleted any residual ISCs by FACS^51^, and assessed CDX2 binding by CUT&RUN-seq. Relative to fetal and homeostatic cell states (ISC, Diff 1, Diff 2; Figure 4A), D5 TA cells showed a broadly reversed pattern: they gained CDX2 binding at fetal- and ISC-specific loci while losing binding at TA-specific loci (Diff 1, Diff 2; Figure 4E). Specifically, 31% of fetal-specific and 56% of ISC-specific loci showed at least a 1.5-fold gain in CDX2 binding in D5 TA cells relative to homeostatic TA cells, while 56% and 62% of Diff 1- and Diff 2-specific loci, respectively, showed at least a 1.5-fold loss (Figure 4F, G, S4B-C). mRNA-seq from D5 TA cells showed a significant increase in expression of genes near fetal- and ISC-specific CDX2-rebound loci (within 25 kb), approaching homeostatic ISC expression levels, while genes bound by CDX2 in differentiated cells (Diff 1, Diff 2) showed significant expression loss (Figure 4H).

Native ChIP-seq for H3K27ac showed significant reactivation of fetal enhancers following CDX2 rebinding after prolonged inactivity spanning fetal development and hundreds of cumulative divisions across the expanding epithelial progeny^52–54^, providing a mechanistic basis for CDX2-driven reactivation of fetal genes (Figure S4D). ISC enhancers, by contrast, retained H3K27ac signal in TA cells, consistent with their more recent activity just a few cell divisions earlier, potentially facilitating CDX2 rebinding during regeneration. Together, CDX2 binding in fetal, homeostatic ISC, and TA cells formed distinct signatures, with the TA-cell signature shifting significantly toward both the fetal and ISC states during regeneration (Figure 4I).

Together, these results show that CDX2 tailors its molecular activity to the situational demands of intestinal epithelial cells: establishing developmental programs during fetal patterning, maintaining differentiation programs during homeostatic ISC differentiation, and, during regeneration, simultaneously redeploying both programs in non-stem cells to reinstate a stem-cell state. CDX2 thus functions as an on-demand regulator of epithelial cell fate, capable of engaging enhancer and gene programs from strikingly distinct temporal origins: fetal programs dormant for hundreds of cell divisions, and ISC programs active only a few cell divisions earlier (Figure 4J).

### CDX2 reengages fetal enhancers and its partner-TF (CTCF) co-occupancy signature during regeneration

Given CDX2’s region-specific engagement with fetal genes during development (Figure 2E-G, 3E) and its rebinding to fetal enhancers in regenerating duodenal TA cells (Figure 4A-B), we asked whether this regenerative reengagement is similarly region-specific, and what molecular features underlie its function in non-stem crypt cells. We determined CDX2 binding in non-stem colonic crypt cells 5 days post-radiation by CUT&RUN (Figure S5A), following ISC depletion by Lgr5-EGFP-based FACS^51^, and compared this to CDX2 rebinding in the duodenum (Figure 4A). CDX2 rebinding at many loci reflected region-specific fetal CDX2 binding, while loci commonly bound in both fetal midgut and hindgut also showed rebinding in both regenerating duodenum and colon (Figure 5A-C). This dual pattern recapitulates both the region-specific and commonly shared components of CDX2’s developmental binding (Figures 1-3), suggesting that CDX2 rebinding during regeneration faithfully mirrors the regional architecture established during fetal development, rather than reactivating a generic, non-specific fetal state.

**Figure 5:**
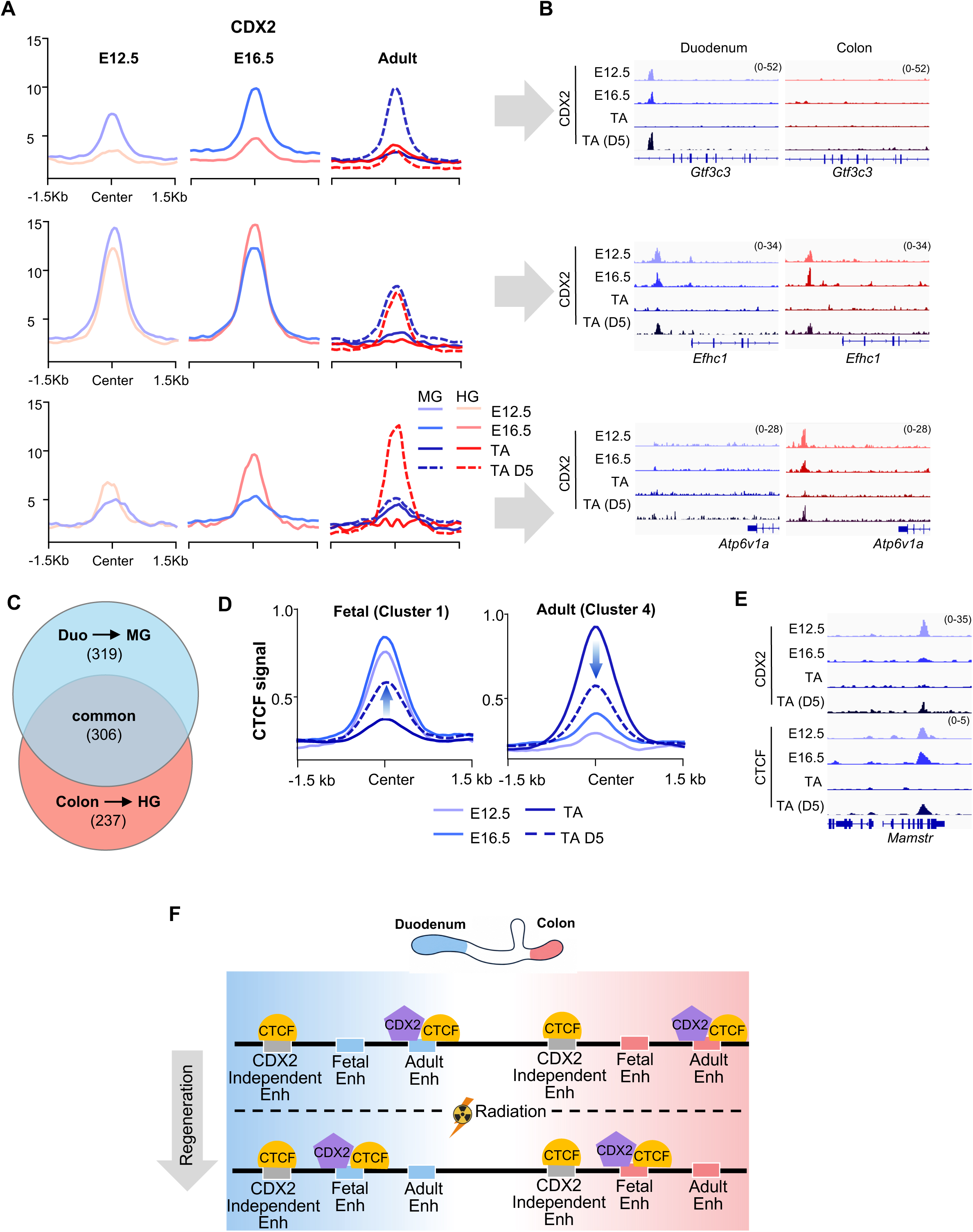
Regions specific rebinding of CDX2 to fetal and ISC enhancers during epithelial regeneration. (A) Aggregate CDX2 binding profiles showing region-specific re-occupancy of fetal-specific CDX2-bound loci during regeneration. CDX2 occupancy is shown in fetal epithelium (E12.5 and E16.5), adult transit-amplifying (TA) epithelium, and day 5 regenerating TA epithelium (TA D5). (B) Representative genome-browser tracks of CDX2 signal illustrating CDX2 binding dynamics at fetal enhancer loci from the clusters shown in panel A. These examples demonstrate selective rebinding of CDX2 to fetal regulatory elements during epithelial regeneration. (C) Venn diagram showing the overlap of regeneration-associated CDX2 rebinding to fetal-specific CDX2-bound loci in the duodenum (duodenum → MG) and colon (colon → HG). A substantial fraction of fetal-specific enhancers shared between the midgut (MG) and hindgut (HG) are rebound by CDX2 during regeneration, whereas others exhibit region-specific rebinding. (D) Average CTCF occupancy centered on fetal cell-specific CDX2 binding (Cluster 1, Fig. 4A) and adult cell-specific CDX2 binding (Cluster 4, Fig. 4A) across development (E12.5, E16.5), adult homeostasis (TA), and regeneration (TA D5). CTCF co-occupancy is dynamically remodeled together with CDX2, reflecting reuse of enhancer activity and chromatin organization associated with fetal developmental during regeneration. (E) Representative genome-browser views of CDX2 and CTCF signal showing coordinated CDX2 and CTCF occupancy at fetal and adult regulatory regions during epithelial regeneration. (F) Model illustrating region-specific rebinding of CDX2 during epithelial regeneration and co-occupancy of CTCF.

As before (Figure 3), we asked whether CDX2’s functional relationship with partner transcription factors was similarly reengaged during regeneration: CUT&RUN-seq for CTCF in regenerating (D5) TA cells showed that loci gaining CDX2 binding at fetal enhancers also gained CTCF co-occupancy, while CTCF binding was reduced at loci retaining differentiated-cell-specific CDX2 binding (Figure 5D-E). Together, these results show that CDX2 reengages fetal enhancer architecture during regeneration with regional and multi-enhancer fidelity, alongside a corresponding CTCF co-occupancy signature (Figure 5F) collectively supporting reconstruction of a hybrid fetal-ISC enhancer network and cell state.

### Selective reactivation of a shared fetal tissue-structural program across diverse injury types is linked to CDX2 re-engagement of fetal binding sites

In vivo injury models — including radiation, toxin exposure, and helminth infection that induce ISC loss — as well as ex vivo organoid systems that impose structural stress on the crypt, each trigger activation of distinct groups of fetal-associated genes^2^. However, several fundamental questions remain unresolved: (a) what general processes reflect cellular and tissue demands in response to ISC loss; (b) whether fetal gene reactivation reflects broad developmental reprogramming or the selective re-engagement of specific processes required to restore tissue integrity and function; (c) the molecular and epigenetic mechanisms underlying this reactivation; and (d) how cells balance stem-state versus tissue-structural demands.

To address this, we first compiled all fetal genes reported as reactivated across intestinal epithelial states, exposures, and injuries in six recent studies contributing seven damage-response gene sets in total (bulk and single-cell RNA-seq^38,55–59^, then determined CDX2 binding these genes (TSS ± 25 kb) across fetal, adult, and regenerating (5 days post-radiation) states. Of 721 unique fetal genes reported to be reactivated during regeneration across these studies, 436 (60%) showed CDX2 binding during development and regeneration (Figure S6A), which we designate the *436-gene CDX2-bound fetal regeneration* panel (Table S1). Within individual published damage-response gene sets, up to 86% of genes showed increased CDX2 binding in regenerating TA cells relative to homeostatic TA cells (minimum 47%; Figure 6A), indicating broad CDX2 involvement in fetal gene reactivation and confirming that this panel holds consistently across diverse injury types, independent of the specific type of tissue damage.

**Figure 6.**
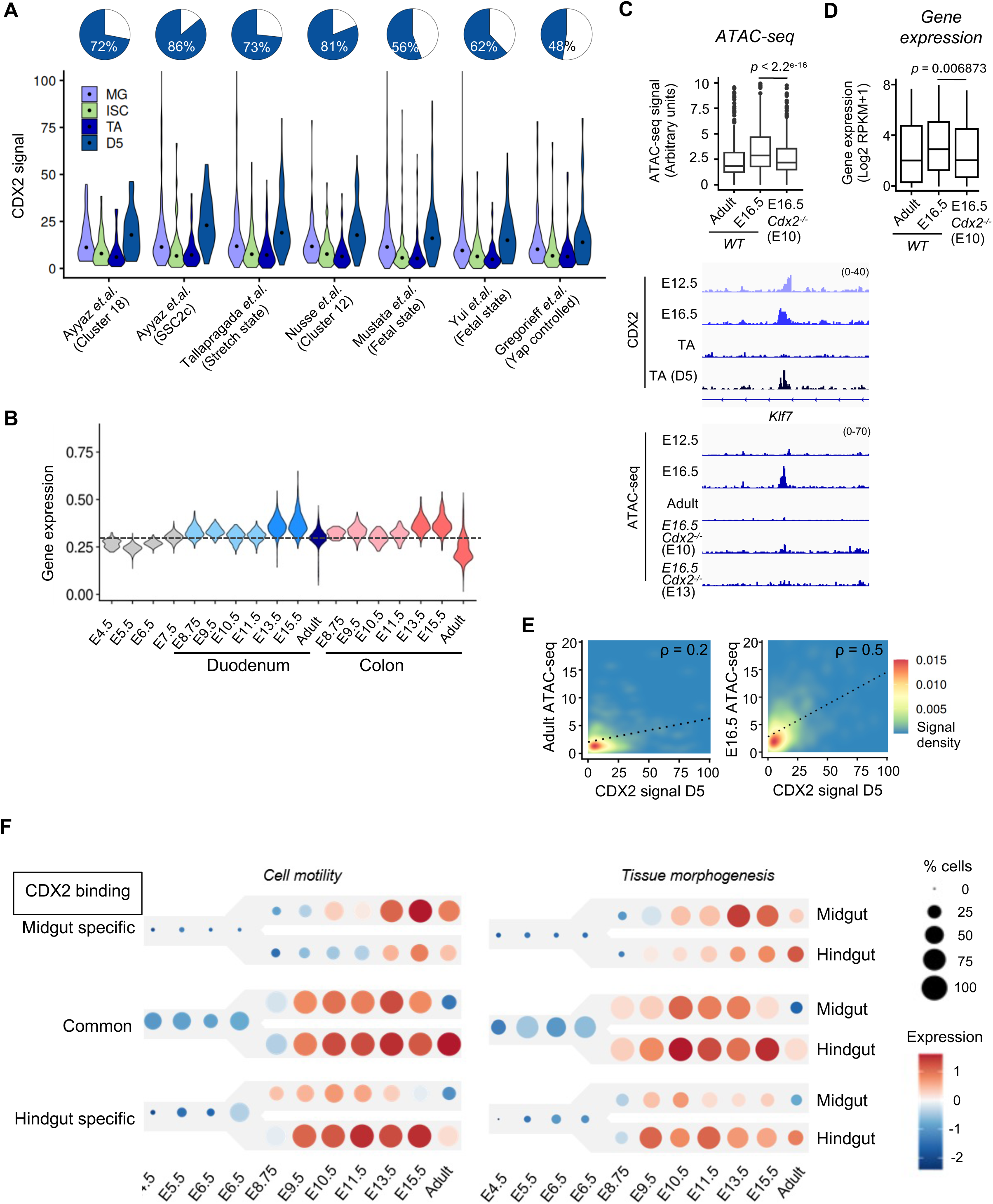
Fetal cell-motility and tissue-morphogenesis programs are reactivated through CDX2 during regeneration. (A) CDX2 occupancy at genes (within ±25 kb of the transcription start site) in various fetal-like gene groups identified in published studies ^38,55–59^. Violin plots show normalized CDX2 CUT&RUN signal across fetal midgut (MG, highest signal between E12.5 and E16.5), adult intestinal stem cells (ISC), transit-amplifying (TA) cells, and day 5 regenerating epithelium (TA D5) for gene groups in various studies. Pie charts above each of the gene groups indicate percentage of genes in each group bound by CDX2. (B) Violin plots showing expression (scRNA-seq) of all unique CDX2 associated genes (n=436) among various published fetal-like gene groups across fetal endoderm development followed by MG → Duodenum (blue) and HG → Colon (red) development and in the duodenum and colon, highlighting the temporal activation and maintenance of region-specific gene expression. Dotted line indicates median expression at E7.5. (C) Box plots (top) showing chromatin accessibility (ATAC-seq) signal at CDX2-bound regulatory elements near (TSS +/- 25 kb) all CDX2 associated fetal genes (n=436, as in B) in wild-type adult intestinal cells (TA cells), fetal intestine (E16.5), and *Cdx2*-deficient fetal intestine (E16.5, *Cdx2*−/−, deletion induced at E10). Representative genome-browser tracks (bottom) of ATAC and CDX2 signal illustrating that loss of CDX2 binding at fetal-development-bound regions in adult cells reduces accessibility at these regulatory elements, and that CDX2 deletion at E10 or E13 similarly causes loss of accessibility at E16.6, indicating that CDX2 is required to establish or maintain an open chromatin state.^12^ (D) Expression of genes associated with fetal CDX2-bound enhancers (as in B) in wild-type adult intestinal cells (TA cells), fetal intestine (E16.5), and *Cdx2*-deficient fetal intestine (E16.5, *Cdx2*−/−, deletion induced at E10). Boxplots show normalized gene expression (log2[RPKM + 1]), demonstrating reduced expression following loss of CDX2. (E) Correlation between CDX2 occupancy during regeneration (day 5) and chromatin accessibility in adult and fetal (E16.5) intestine. Density plots compare CDX2 binding with adult (left) or fetal (E16.5; right) ATAC-seq signal, illustrating preferential reactivation of developmentally accessible regulatory elements during regeneration. (F) Dot plots showing expression (scRNA-seq) of cell motility and tissue morphogenesis genes with fetal cell specific CDX2 occupancy in midgut, hindgut or both (common), during fetal development and adult cell states of MG → Duodenum and HG → Colon lineages. These data demonstrate that CDX2 engages region-specific fetal enhancers to activate temporally fetal cell- and region-specific as well as commonly used migratory and morphogenetic gene programs.

To determine the developmental window during which this panel is active, prior to its inactivation in adult epithelium, we used integrated scRNA-seq data spanning 10 developmental timepoints (Figure 2A). Consistent with rising CDX2 expression after E8 (Figure 2A), the *436-gene CDX2-bound fetal regeneration* panel was upregulated by E8.75 in both duodenal and colonic endodermal precursors relative to the earlier epiblast (E4.5– E6.5) and definitive endoderm (E7.5) stages, then declined in both adult duodenal and colonic epithelia (Figure 6B), suggesting that CDX2-bound fetal regeneration genes broadly support processes important for intestinal epithelial development across regions. Consistent with CDX2’s direct involvement in this fetal program, chromatin accessibility (ATAC-seq) at CDX2-bound enhancers near these genes was higher at E16.5 than in adult TA cells, and *Cdx2* deletion at E10 significantly reduced this accessibility in fetal tissue (Figure 6C). Expression of the linked genes was similarly reduced following *Cdx2* deletion (Figure 6D).

Moreover, CDX2 binding at the enhancers of this panel in post-injury crypt cells correlated more strongly with fetal (E16.5) than with homeostatic chromatin accessibility (Figure 6E, S6B), consistent with CDX2 reactivating a fetal regulatory program during regeneration rather than simply binding generically accessible chromatin. Together with its damage-type-independent origin (Figure 6A) and its developmental expression trajectory (Figure 6B), the *436-gene CDX2-bound fetal regeneration* panel links this gene set to a complete CDX2 occupancy trajectory spanning fetal, adult, and regenerating epithelial states, extending prior expression-based fetal-reprogramming signatures with direct evidence of the transcription factor associated with their reactivation. To determine which fetal processes are being reinstated, we performed gene ontology enrichment on this panel and found significant enrichment for cell motility and tissue morphogenesis (Figure S6C), concordant with our earlier finding that CDX2’s motility- and morphogenesis-related targets are predominantly genes commonly activated in both midgut and hindgut during development (Figure 2E-G, S2E). Reinforcing this pattern, gene ontology analysis of the CDX2 loci commonly rebound across regenerating duodenum and colon (Figure 5A) were also enriched for tissue-structural processes, while region-specific rebound loci were instead enriched for cell-proliferation and growth-factor-response genes (Figure S6D). This convergent enrichment across two independent gene sets suggested that CDX2 selectively borrows a specific subset of fetal processes to support regeneration.

To test this directly, we identified all CDX2-bound cell-motility and tissue-morphogenesis genes within the comprehensive set of CDX2 loci commonly and distinctly bound across midgut and hindgut development (Figure 2E-G, S2E), the most complete developmental CDX2-binding dataset available, yielding 314 cell-motility and 517 tissue-morphogenesis genes, which we designate the *CDX2-bound fetal cell-motility signature* and the *CDX2-bound fetal tissue-morphogenesis signature*, respectively (Table S2). We tracked their activity across early endoderm development into the fetal midgut and hindgut lineages using integrated scRNA-seq data (Figure 2A, 6F). Region-specific targets were activated beginning around E7.5 and subsequently downregulated in adult cells. Genes co-bound by CDX2 in both lineages, however, were already active before regional diversification and before CDX2 expression itself began at E7.5 (Figure 2A), indicating that these loci are first engaged by other transcription factors or other activating mechanisms and later amplified by CDX2 binding (Figure 6F). Combined with the CDX2-dependence of these genes during development (Figure 6C) and their reactivation across multiple injury models (radiation, helminth infection, and organoid stretch) through CDX2 rebinding, these results show that CDX2 establishes region-specific tissue-morphogenesis programs during development and selectively reengages them during regeneration. Cell migration, cell-cell adhesion, and tissue-structural programs thus evolve as the midgut and hindgut give rise to the small intestine and colon, with CDX2 acting as a central regulator: establishing these region-specific programs during development, and reengaging both region-specific and shared fetal motility/morphogenesis programs precisely when regeneration demands them. This tissue-structural reactivation still represents only one demand facing regenerating cells, which must simultaneously reestablish the stem-cell compartment itself — raising the question of how, and in which cells, CDX2 balances these structural and stem-cell programs during regeneration.

### Distinct spatial deployment of fetal tissue-structural and stem-cell programs during intestinal regeneration is linked to temporally divergent CDX2 re-occupancy

CDX2 reengages the fetal tissue-structural programs captured by the CDX2-bound fetal cell-motility and tissue-morphogenesis signatures during regeneration (Figure 6), yet regenerating cells face a second, parallel demand: reestablishing the stem-cell compartment itself. Whether individual cells meet both demands simultaneously, or whether these fetal signatures and the stem-cell program are distributed across distinct cell populations along the crypt-villus axis, remains unclear.

To address this, we performed single-cell multiomic analysis (scRNA-seq and scATAC-seq) on regenerating crypts, analyzing 4,642 homeostatic and 4,458 regenerating (D4.5 post-radiation) cells after quality control. Joint dimensionality reduction (UMAP) of both modalities (Figure S7A) identified ISC, TA1, TA2 (non-stem crypt clusters proximal and distal to the ISC state, respectively), early enterocyte, enterocyte, and secretory populations in homeostatic tissue using established markers (Figure 7A, S7B–C). Regenerating cells showed substantially greater mixing among crypt populations (Stem, TA1, TA2) than the well-separated clusters seen at homeostasis (Figure 7B), demonstrating that multiome profiling captures cell-state dynamics during regeneration. To model the continuous ISC→TA→enterocyte differentiation trajectory, we performed pseudotime analysis (Monocle^60^) using the 500 most variable genes, including genes linked to dynamic CDX2 binding across fetal, ISC, and differentiated states during development and regeneration (Figure 4A), establishing an ISC differentiation trajectory along which cells could be ordered (Figure 7C).

**Figure 7.**
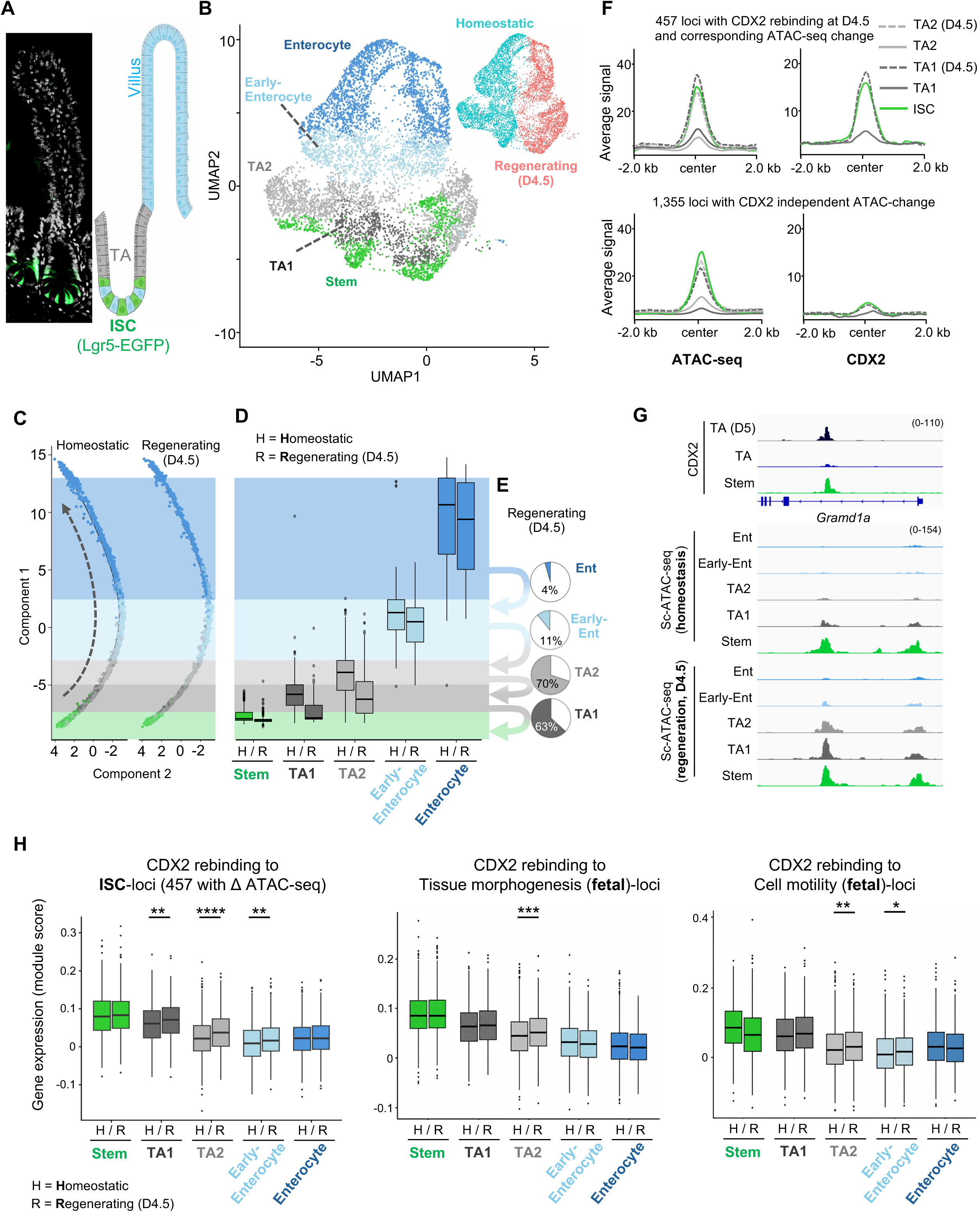
TA cells exhibit a CDX2-driven dual fetal/stemness signature during intestinal epithelial regeneration. (A) Representative fluorescent micrograph showing crypt-villus structure within the mouse intestinal epithelium, with Lgr5+ ISCs at the bottom of the crypt showing endogenous EGFP signal; schematic (right) shows ISC, transit-amplifying (TA), and villus cells. (B) UMAP of scMultiome (scRNA-seq and scATAC-seq) from intestinal epithelium showing cell states along the ISC → Enterocyte differentiation path in homeostatic (uninjured) and regenerating epithelium (day 4.5 post-radiation); the inset (top-right) identifies homeostatic (cyan) and regenerating (pink) cell populations. (C) Homeostatic and regenerating (D4.5) cells arranged on pseudotime projection generated using top 500 most differential features including CDX2 linked genes (Figure 4A); dotted arrow indicates the ISC → Enterocyte differentiation trajectory in the homeostatic cells. (D) Box and whisker plots showing distribution for each of the five cell states—stem, TA1, TA2, early enterocyte, and enterocyte—in homeostatic and regenerating (D4.5) cells; bottom and the top of the box represent end of the first and the third quartile and the line within the box represents end of second quartile. (E) The pie charts show the proportion of cells from each of the five cell states—stem, TA1, TA2, early enterocyte, and enterocyte—that map to the territory of the preceding cell-state along the differentiation trajectory; cell state territories are defined as the territory between the third quartile of a cell state and the third quartile of the cell state before that within the homeostatic trajectory, as indicated by the light shaded bands with colors corresponding to each of the cell state across C and D); e.g., 63% of regenerating TA1 cells map to third quartile or lower within the homeostatic stem-cell state. Compared with WT cells, regenerating (D4) cells were more frequently assigned to earlier states, revealing increased representation of stem-like cells and immature TA1, TA2, and early enterocyte populations. This shift toward a less differentiated phenotype was not observed among mature enterocytes. (F) Average ATAC-seq (left) and CDX2 (right) signal profiles centered on two sets of loci. Top, 457 loci with CDX2 rebinding at D4.5 and a corresponding increase in ATAC-seq signal. Bottom, 1,355 loci with an ATAC-seq signal change at D4.5 that occurs without a corresponding change in CDX2 binding. Signal is shown in homeostatic ISC, TA1, and TA2 populations, and in the corresponding regenerating TA1 (D4.5) and TA2 (D4.5) populations. (G) Representative genome-browser tracks of ATAC and CDX2 signal at a stem-cell gene locus *Grand1a*^77^ showing CDX2 binding and ATAC-seq (sc-ATAC-seq pseudo-bulk) signal lost in TA cells and regained in regenerating (D4.5) TA1 and TA2 cells. (H) Boxplots showing gene expression scores for the ISC signature (457 CDX2-rebound loci with corresponding ATAC-seq change), the fetal tissue-morphogenesis signature, and the fetal cell-motility signature (left to right), in homeostatic (H) and regenerating (R) cells within the five cell states spanning the differentiation trajectory. Center lines indicate medians, boxes indicate interquartile ranges (IQRs), whiskers extend to 1.5 × IQR, and points indicate outliers. Statistical significance was determined using the Wilcoxon rank-sum test (*, *p* < 0.05; **, *p* < 0.01; ***, *p* < 0.001; ****, *p* < 0.0001). Together, these analyses show that CDX2-bound ISC programs are reactivated broadly across regenerating cell states, while CDX2-bound fetal programs are reactivated transiently and in select crypt populations, to restore epithelial structure and function.

To quantify cell-state changes during regeneration, we defined cell territories along this trajectory as the interval between the third quartile of one cell state and the third quartile of the preceding state under homeostatic conditions (Figure 7D), then asked how cells from each of the five states redistributed across these territories during regeneration relative to homeostasis. At D4.5 post-radiation, 63% of TA1 cells mapped to a stem-like territory and 70% of TA2 cells mapped to the TA1 territory, indicating a shift toward a less differentiated state that was prominent among TA populations but sharply diminished in cells further along the differentiation trajectory (early enterocytes and enterocytes) (Figure 7E).

Because CDX2 re-binds a substantial fraction of its ISC-specific target enhancers in regenerating crypt cells (Figure 4A), we next examined accessibility changes in TA populations post-radiation to determine how CDX2 binding dynamics relate to these cell-state shifts. Comparing TA accessibility to homeostatic ISC populations (2-fold change) identified 1,812 loci that rapidly lost accessibility during ISC-to-TA differentiation; these same loci regained accessibility in the corresponding regenerating TA clusters (D4.5-TA1, D4.5-TA2), indicating that ISC-like enhancers are re-engaged in non-stem cells during regeneration. Of these, 457 loci (34%) showed increased

CDX2 binding (Figure 7F,G, S7D), suggesting that CDX2 rebinding reopens stem-specific enhancers silenced during differentiation, potentially alongside engagement of other transcription factors. An additional 1,461 loci showed reduced (>1.5-fold) or lost CDX2 binding during ISC differentiation and were re-bound by CDX2 during regeneration without a corresponding change in accessibility (Figure S7D), suggesting that chromatin at these loci is not fully remodeled during differentiation, allowing CDX2 to rapidly reengage them once regeneration begins. On the other hand, 1,355 loci showed accessibility changes during ISC differentiation that were reversed during regeneration independent of CDX2 binding, suggesting parallel involvement of other transcription factors (Figure 7F). Together, these results indicate that ISC regeneration involves extensive re-engagement of stem-cell enhancers, predominantly — though not exclusively — through selective CDX2 rebinding. CDX2 concurrently reactivates two temporally distant classes of enhancers during regeneration: recently used ISC-specific enhancers (Figure 4A, C), and fetal enhancers from the distant developmental past, reengaged after hundreds of intervening cell divisions^52–54^ to activate the CDX2-bound fetal cell-motility and tissue-morphogenesis signatures (Figure S2B). To determine how CDX2 balances this concurrent reactivation, we asked which cell states along the differentiation trajectory activate each program. CDX2-bound ISC-linked genes that quickly lost CDX2 binding and ATAC-seq signal in TA cells showed significant reactivation broadly, across the entire crypt (TA1, TA2) and early enterocyte populations, during regeneration (Figure 7G,F). Genes linked to enhancers with chromatin accessibility changes independent of CDX2 binding showed a similarly broad increase in expression, across regenerating stem, TA1, and TA2 populations (Figure S7E). In contrast, genes within the CDX2-bound fetal tissue-morphogenesis signature were significantly and selectively activated in the TA2 cluster alone, whereas genes within the CDX2-bound fetal cell-motility signature were activated in both TA2 and early enterocyte clusters (Figure 7F). CDX2 thus draws simultaneously on two very different enhancer histories: ISC-specific enhancers used only recently, and fetal enhancers untouched since early development, a gap that cell division and differentiation would ordinarily be expected to erase. Despite this, CDX2 deploys each transiently, but with distinct spatial reach: ISC programs are reactivated broadly, across multiple cell tiers in the regenerating epithelium, while fetal programs are reactivated in select crypt populations. This dual capacity underscores CDX2’s role as an on-demand, cell-state-specific regulator during tissue repair. Together with the region-specific, developmentally established CDX2 binding patterns described earlier, these findings show that regeneration does not broadly reinstate a fetal state, but selectively and precisely redeploys the same spatially and temporally organized regulatory framework CDX2 first established during development.

## DISCUSSION

Cell motility and tissue morphogenesis are essential for establishing tissue structure. In barrier epithelia such as the intestine, tight cell-cell adhesion allows a single cell layer to separate underlying tissue from the harsh luminal environment, and loss of this barrier integrity contributes to a wide range of diseases^61,62^. Epithelial tissues have therefore evolved not only to establish correct structure during development, but to rapidly and precisely reactivate motility and morphogenesis genes following injury, enabling swift barrier repair and a return to normal structure and function^63^.

Recent studies show that many fetal genes, including those governing motility and structure, are reactivated during intestinal regeneration^2^, raising the question of whether this reflects broad reversion to a fetal-like state or a more selective process. While these studies show that fetal-like programs are reactivated during regeneration; here, we define which programs are selectively reengaged, how they are organized within a CDX2-directed enhancer framework, and why their coordinated reactivation enables faithful, region-specific restoration of tissue structure and function.

We show that these programs — defined here as the *CDX2-bound fetal cell-motility signature* and the *CDX2-bound fetal tissue-morphogenesis signature* — are activated early in development, before regionalization, then become selectively amplified, broadly or region-specifically, before their activity is minimized in adult tissue. Following regional specification, CDX2 directly binds many of these genes, with a marked increase in expression, particularly among motility and morphogenesis genes shared between duodenum and colon — suggesting that, in the late fetal window preceding birth, CDX2 binding and enhancer activation prepare the tissue’s structural architecture for postnatal function. In adult tissue, CDX2 no longer occupies these enhancers, and their expression is markedly reduced.

Following epithelial injury or disease states associated with stem cell loss, the epithelium must simultaneously recover lost stem cells and rebuild tissue structure. The intestine is known to increase cell motility to repair barrier breaches and replace lost stem cells, and Wnt, Bmp, Notch, and Yap signaling, which are critical for fetal epithelial development, have been implicated in these processes^2,38,64–73^. However, the central transcription factors that directly activate the genes required for this recovery have remained largely undefined.

By compiling gene sets activated across multiple independent injury models and comparing them with midgut and hindgut developmental expression dynamics, we assembled the *436-gene CDX2-bound fetal regeneration panel* — a cross-study, CDX2-occupancy-defined resource spanning fetal, adult, and regenerating states. This reveals that, alongside stem-cell gene reactivation, the CDX2-bound fetal cell-motility and tissue-morphogenesis signatures represent a major, previously undefined component of the fetal programs reengaged during regeneration, and that these genes are direct CDX2 targets in both development and regeneration.

Several crypt populations beyond the stem compartment, including +4 cells above the stem zone and isthmus cells at the upper crypt, contribute to ISC regeneration and epithelial maintenance^74–76^, though how these positions are recruited into a regenerative response has remained unclear. Using CDX2 binding profiles across homeostatic and regenerating crypt populations together with single-cell multiome analysis, we find that activation of the CDX2-bound fetal signatures varies systematically along the crypt axis and tracks each population’s transition toward a stem-like state. TA2 cells, positioned farthest from the stem compartment, analogous to the isthmus, show the strongest activation of these fetal signatures, suggesting CDX2 may reprogram precisely these distal, normally non-regenerative populations to contribute to repair. CDX2-bound ISC-linked genes, by contrast, are activated broadly across the entire crypt, indicating a more spatially permissive mode of reactivation than the region-restricted fetal signatures.

As master regulator of intestinal lineage, CDX2 is essential for intestinal development; its loss abolishes intestinal tissue formation. Its roles in adult stem cell differentiation into secretory and absorptive lineages, and in the function of these cells, are well established^10–15,41^. However, CDX2’s direct involvement in region-specific gene activation across development, adult tissue, and regeneration has remained underappreciated. We therefore profiled CDX2 binding across ten cellular states spanning development, homeostasis, and regeneration, including midgut and hindgut development, their adult derivatives (duodenum, ileum, colon), and ISC and crypt populations under homeostatic and post-injury conditions.

GATA4 and PDX1 mark small intestinal identity and SATB2 marks colonic identity, yet what drives their own region-specific expression has been unclear. We show that CDX2, despite uniform expression across midgut and hindgut progenitors, precedes these regulators in expression, directly occupies their genes, and binds their region-specific enhancers, while also cooperating with CTCF at these loci, potentially facilitating regional chromatin organization. Together, these findings describe a hierarchical transcription factor network in which CDX2 both regulates downstream regional TFs directly and facilitates partner-factor recruitment to regulatory elements. In adult tissue, CDX2 binding remains region-specific, with enhancer number tracking regional expression level, while stem-cell and differentiation genes remain CDX2-bound across both regions, indicating CDX2 functions simultaneously as a region-specific and broadly shared regulator.

Together, CDX2’s developmental and regenerative roles are mechanistically interconnected. During fetal development, CDX2 directly targets the CDX2-bound fetal cell-motility and tissue-morphogenesis signatures, amplifying their expression; in adult tissue these enhancers are no longer CDX2-bound and expression is minimized, providing a mechanistic basis for the tissue agenesis previously observed upon CDX2 loss. During regeneration, CDX2 reengages these same fetal enhancers, potentially facilitating cellular mobilization and re-establishment of epithelial architecture in a manner recapitulating development. Although these signatures are largely shared between small intestine and colon, CDX2 additionally supports region-specific developmental programs, including differentiation, regional signaling, and adult function, while reactivating these programs simultaneously alongside ISC-specific enhancers in non-stem transit-amplifying cells.

This reveals an unexpected feature of CDX2-mediated regeneration: simultaneous engagement of two temporally distinct regulatory reservoirs. CDX2 binds fetal enhancers that facilitated development hundreds of cell divisions earlier, while re-engaging ISC-specific enhancers occupied only a few cell divisions previously^52–54^ — all while preserving regional fidelity. Regenerative reprogramming is therefore not merely a reversal of differentiation, but a highly selective process in which CDX2 retrieves and combines programs separated by vastly different developmental distances to reconstruct adult epithelial identity and function, consistent with its role as an on-demand, cell-state-specific regulator of epithelial repair.

More broadly, these findings establish that CDX2 retains the capacity to retrieve and redeploy developmental regulatory information long after organogenesis is complete — engaging stemness, motility, and morphogenesis programs precisely and transiently, in the specific cell types where they are needed, rather than allowing tissue-wide or sustained activation. This capacity for rapid, coordinated, yet tightly restrained access is likely essential not only for injury repair but for any context demanding fast tissue remodeling. Notably, these same programs are hallmarks of malignant growth, raising the possibility that CDX2’s precise, on-demand control may be co-opted in disease states such as cancer, where the same programs become persistently, rather than transiently, activated.

## METHOD DETAILS

### Mouse Models

All animal experiments were conducted under protocols approved by the Institutional Animal Care and Use Committee (IACUC) at the University of Southern California. Mice were maintained under standard housing conditions at 23 ± 1 °C and 50 ± 10% relative humidity on a 12-hour light/dark cycle. Animals exhibiting signs of distress, including hunched posture, ruffled fur, decreased activity, or rapid body weight loss of ≥20%, were removed from the study and euthanized humanely. Euthanasia was carried out by CO₂ inhalation for 5 minutes (∼20% chamber volume/min), followed by cervical dislocation as a secondary method, in accordance with the AVMA Guidelines for the Euthanasia of Animals (2013). Embryos obtained by cesarean delivery were euthanized by decapitation. ROSA26^LSL-tdTomato^ mice were purchased from The Jackson Laboratory. Eed^fl/fl^, Lgr5^EGFP-IRES-DTR^, and Villin^CreER-T^^2^ mouse lines were generously provided by S. Orkin (Boston Children’s Hospital), F. de Sauvage (Genentech), and S. Robine (Institut Pasteur), respectively. Genotypes of all transgenic mice were verified by PCR at weaning and reconfirmed throughout the course of the experiments. To induce Eed deletion, adult mice (≥8 weeks old) received intraperitoneal injections of tamoxifen (2 mg/day) for five consecutive days. Tissue samples were harvested at the indicated experimental time points.

### Isolation of mouse intestinal epithelial cells from E12 and E16 embryos and adult duodenum, ileum, and colon

Embryonic intestinal epithelial cells and adult duodenal and ileal epithelial populations, including villus, intestinal stem cell (ISC), and transit-amplifying (TA) cells, were isolated as previously described (PMID:39843425). Briefly, villus epithelial cells were isolated from the proximal one-third of the small intestine of adult (≥8-week-old) male and female mice. Intestinal tissue was opened longitudinally, rinsed with ice-cold PBS, and incubated in 5 mM EDTA/PBS (pH 8.0) at 4 °C for 30 minutes with gentle agitation. Detached epithelial fractions were passed through a 70-µm cell strainer, and villi were collected in PBS, pelleted by centrifugation (300 × g, 5 minutes), and dissociated in 4% TrypLE prepared in DMEM for 30 minutes at 37 °C. Digestion was quenched with DMEM, followed by cell pelleting. For embryonic epithelial cell isolation (E12.5 and E16.5), intestines from four embryos were dissected, finely minced, and digested in 0.25% trypsin at 37 °C for 20 minutes. Enzymatic digestion was terminated by adding DMEM supplemented with 2% FBS. Cells were centrifuged (500 × g, 5 minutes, 4 °C), resuspended in PBS containing 2% BSA, and EpCAM-positive epithelial cells were enriched using the EasySep™ system with an anti-EpCAM antibody (Thermo Fisher, 13-5791-82). For isolation of ISCs and TA cells, proximal intestinal tissue from Lgr5^EGFP-Cre^ and Lgr5^EGFP-IRES-DTR^ mice, respectively, was processed using the same epithelial dissociation protocol, except tissues were incubated in 5 mM EDTA/PBS (pH 8.0) at 4 °C for 40 minutes before filtration through a 70-µm strainer. Isolated crypts were dissociated in 4% TrypLE for 30 minutes at 37 °C. Viable EpCAM-positive cells (Thermo Fisher, 48-5791-82) were then sorted on a BD SORP FACSymphony S6 to isolate EGFP^hi intestinal stem cells and EGFP^- transit-amplifying cells.

### Adult colon epithelial cells

Following tissue collection, colons were flushed with ice-cold Hank’s Balanced Salt Solution (HBSS), opened longitudinally, and cut into small pieces. Tissue fragments were transferred to 50-mL conical tubes containing 15 mL of epithelial dissociation solution consisting of HBSS supplemented with 100 U/mL penicillin/streptomycin, 2% fetal bovine serum, 100 μg/mL DNase I, 10 mM HEPES, and 10 mM EDTA. Samples were incubated at 37 °C with gentle agitation every 5 minutes for a total of 15 minutes, followed by cooling on ice for 10 minutes. Tissues were then transferred to fresh HBSS and shaken vigorously for 45 seconds to release colonic crypts. Crypt fractions were collected by centrifugation and resuspended in 5 mL of prewarmed TrypLE containing 100 μg/mL DNase I. After incubation at 37 °C for 5 minutes, samples were gently triturated until a single-cell suspension was obtained, as confirmed microscopically. Digestion was terminated by adding 1 mL of fetal bovine serum. Cells were passed through a 40-μm cell strainer, washed, and processed for downstream applications. EpCAM-positive epithelial cells were subsequently enriched using the EasySep™ system with an anti-EpCAM antibody.

### Irradiation of mouse intestine and colon

Mice were exposed to irradiation in accordance with approved IACUC protocols. Immediately before irradiation, animals were anesthetized by intraperitoneal injection of ketamine (MWI Animal Health, Boise, ID) and xylazine (Anased LA, 100 mg/mL; USC Department of Animal Resources, Los Angeles, CA), diluted in sterile PBS and administered at doses of 60 mg/kg and 8 mg/kg, respectively. Ophthalmic ointment was applied to prevent corneal drying during the procedure. Mice received a single dose of 10 Gy X-ray irradiation using an X-RAD 320 irradiator (Precision X-Ray, Madison, CT) at a dose rate of 2.0 Gy/min. Irradiation was restricted to the abdominal region, while the remainder of the body was protected with lead shielding. Following irradiation, animals were returned to clean cages containing fresh bedding, food pellets placed on the cage floor, HydroGel (ClearH2O, Westbrook, ME), and water available ad libitum. Mice were closely monitored until they had fully recovered from anesthesia.

### Whole-intestine endogenous fluorescence imaging

Intestinal tissues were dissected, fixed overnight in 4% paraformaldehyde at 4 °C, and washed five times with PBS. Tissues were cryoprotected by incubation in 30% sucrose prepared in PBS overnight at 4 °C, embedded in optimal cutting temperature (OCT) compound, and stored at −80 °C until processing. Cryosections (7 μm) were generated using a Leica CM3050 S cryostat and mounted with VECTASHIELD Antifade Mounting Medium containing DAPI (Vector Laboratories, H-1200-10) for nuclear counterstaining. Endogenous fluorescence was used to distinguish Lgr5^+^ cell populations, which were visualized using the FITC channel, respectively. Entire intestinal sections were imaged with a Zeiss Axioscan.Z1 slide scanner.

### Native ChIP–seq

Native ChIP–seq was carried out as previously described. Briefly, cells were lysed in buffer containing 0.1% Triton X-100, 0.1% sodium deoxycholate, and protease inhibitor cocktail. Chromatin was digested with micrococcal nuclease (MNase; New England Biolabs, M0247S) at room temperature for 5 minutes, and the reaction was terminated by adding 5 μL of 0.25 mM EDTA. Digested chromatin was pre-cleared by incubation with anti-IgA magnetic Dynabeads (Thermo Fisher, 10001D) for 2 hours, followed by overnight immunoprecipitation at 4 °C using antibody-bead complexes containing 0.5 μg of anti-H3K27me3 or anti-H3K27ac antibodies (Diagenode, C15410195 and C15410196) in immunoprecipitation buffer (20 mM Tris-HCl, pH 7.5, 2 mM EDTA, 150 mM NaCl, 0.1% Triton X-100, and 0.1% sodium deoxycholate). Immunoprecipitates were washed twice each with low-salt wash buffer (20 mM Tris-HCl, pH 8.0, 2 mM EDTA, 150 mM NaCl, 1% Triton X-100, 0.1% SDS) and high-salt wash buffer (20 mM Tris-HCl, pH 8.0, 2 mM EDTA, 500 mM NaCl, 1% Triton X-100, 0.1% SDS). DNA was eluted in elution buffer (1% SDS, 100 mM sodium bicarbonate) for 1.5 hours at 65 °C. Histone proteins were digested with protease (Qiagen, 19155) for 30 minutes at 50 °C, and DNA fragments were purified using Sera-Mag magnetic beads in 30% PEG. Sequencing libraries were prepared as previously described by end repair, 3′ A-tailing, and ligation of Illumina sequencing adapters (New England Biolabs, E6000B-10). Libraries were indexed, amplified by PCR for 10 cycles, and sequenced on an Illumina NovaSeq 6000 platform at Genewiz (Azenta Life Sciences).

### CUT&RUN

CUT&RUN assays were performed using the ChIC/CUT&RUN Assay Kit (Active Motif, 53180) according to the manufacturer’s instructions with minor modifications. Freshly isolated epithelial cells were resuspended in permeabilization buffer consisting of ChIC/CUT&RUN Wash Buffer supplemented with 0.01% digitonin and 1× protease inhibitor cocktail and incubated on ice for 5 minutes. Cells were subsequently immobilized on Concanavalin A–coated magnetic beads by incubation at room temperature for 10 minutes. Bead-bound cells were resuspended in 180 μL of antibody buffer (permeabilization buffer containing 2 mM EDTA) supplemented with 0.5 μg of either anti-Cdx2 (Cell Signaling, D11D10) or anti-CTCF (Diagenode, C15410210) antibody and incubated overnight at 4 °C. Following antibody binding, MNase was recruited to the nuclei by incubation at 4 °C for 1 hour, after which chromatin digestion was carried out for 30 minutes at 4 °C on a nutator. Cleaved DNA fragments were released by incubation at 4 °C for 30 minutes and purified using Sera-Mag magnetic beads in 30% PEG-8000. Sequencing libraries were generated as described above for the Native ChIP–seq protocol.

### RNA-seq

Total RNA was isolated using the mirVana™ miRNA Isolation Kit (Thermo Fisher Scientific, AM1561). RNA-seq libraries were generated with the NEBNext® UltraExpress RNA Library Prep Kit (New England Biolabs, E3330S) according to the manufacturer’s instructions. Libraries were sequenced on an Illumina NovaSeq 6000 platform at Genewiz (Azenta Life Sciences) to generate 150-bp paired-end reads.

### Single-cell multiome library preparation (scRNA-seq and scATAC-seq)

Single-cell multiome libraries were generated using the Chromium Next GEM Single Cell Multiome ATAC + Gene Expression kit (10x Genomics) following the manufacturer’s instructions. Briefly, isolated nuclei were subjected to Tn5-mediated transposition, followed by reverse transcription and simultaneous barcoding within individual Gel Beads-in-Emulsion (GEMs). Gene expression and ATAC libraries were amplified and purified independently. Final libraries were quantified using the Qubit dsDNA HS Assay (Thermo Fisher Scientific), and library quality was evaluated with an Agilent Bioanalyzer. Sequencing was carried out on an Illumina NextSeq 6000 platform using paired-end reads with the following configuration: 28 bp for Read 1, 90 bp for Read 2, and a 10-bp i7 index read.

## QUANTIFICATION AND STATSTICAL ANALYSIS

### Computation analyses

#### ChIP-seq and CUT&RUN analysis

Raw ChIP-seq and CUT&RUN sequencing reads were processed using Trim Galore (v0.6.6) to remove adapter sequences and low-quality bases. Filtered reads were aligned to the reference genome using BWA-MEM (v0.7.17) with default settings. Reads that mapped to multiple genomic locations, PCR duplicates, and reads with a mapping quality (MAPQ) score <5 were excluded using SAMtools (v1.17). Peak calling was performed with MACS2 (v2.2.7.1) using a q-value threshold of 0.01, applying narrow peak mode for H3K27ac and transcription factor datasets and broad peak mode for H3K27me3 datasets. Genome-wide coverage tracks (BigWig files) were generated with deepTools (v3.5.1) using a bin size of 10 bp and a smoothing length of 30 bp, followed by quantile normalization with Haystack. Regions overlapping the ENCODE blacklist were removed using BEDTools (v2.30.0).

#### RNA-seq analysis

Raw RNA-seq reads were aligned to the mouse reference genome (GRCm38, GENCODE v25) using STAR aligner (v2.7.8a) in two-pass mode. Sequencing quality was assessed by evaluating per-base sequence quality, per-read GC content (∼50%), strand alignment balance, exon-to-intron read distribution, and 3′ bias. Gene-level read counts were generated using HTSeq (v0.13.5) and normalized across samples with DESeq2. Normalized expression values were subsequently converted to reads per kilobase of transcript per million mapped reads (RPKM). Differential gene expression analysis was performed using DESeq2, with significance determined using the false discovery rate (FDR) thresholds indicated throughout the manuscript. Sequence alignment and all primary processing steps were executed within a Snakemake workflow (v6.6.1).

#### Motif enrichment analysis

Motif enrichment analysis was performed using the *findMotifsGenome.pl* function in HOMER with default parameters, using the full length of each peak or transcription factor binding site as input. Individual motif occurrences were identified as previously described.

### ECDF cutoff based peak selection for histone and TF peaks

Stricter cutoffs for MACS2-called peaks for histone modification and transcription factor CUT&RUN datasets were made using empirical cumulative distribution function (ECDF)-based analysis. For each MACS2-called peak, signal values were extracted from genome-wide normalized bigWig files using deepTools multiBigwigSummary. ECDF curves were calculated for each dataset using the stat_ecdf function from ggplot2. Based on the ECDF distributions, a normalized signal cutoff was selected to identify the transition between background and enriched signal and applied uniformly across datasets. MACS2 peaks with signal values exceeding this threshold were used for downstream analyses.

### Redundant enhancer analysis

Enhancer redundancy was defined as the multiple enhancer regions being assigned to a gene in each biological condition.To quantify enhancer redundancy, MACS2-called peaks across all conditions were combined into a single nonredundant peak set. Promoter-overlapping regions were excluded to restrict the analysis to putative enhancers. Each enhancer region was assigned to the nearest protein-coding gene whose transcription start site (TSS) was located within 25 kb. The ECDF-derived signal threshold was then applied independently to each sample to determine the presence or absence of a peak within each enhancer region. For each gene, the number of associated enhancer regions that passed the threshold was calculated. Gene-specific enhancer counts were then compared between conditions to identify gains and losses in regulatory elements.

### scMultiome data processing, QC

#### Single-cell multiome data analysis

Raw single-cell multiome sequencing data were processed and aligned using Cell Ranger ARC (10x Genomics, v2.0.0). Integrated analysis of single-cell RNA-seq (scRNA-seq) and single-nucleus ATAC-seq (snATAC-seq) datasets was performed using Seurat v4 and Signac. Stringent quality control filters were applied prior to downstream analyses. For the scRNA-seq data, cells were retained if they contained >12,000 unique molecular identifiers (UMIs), >1,000 detected genes, a mitochondrial transcript fraction <0.35, and an expression novelty score >0.8. For the snATAC-seq data, nuclei were retained based on the following criteria: >40% of reads mapping to peaks, >1,000 ATAC fragments per nucleus, blacklist fraction <0.05, transcription start site (TSS) enrichment >2, and nucleosome signal <4. Following quality filtering, 9,100 high-quality cells were retained for downstream analyses, including 4,642 wild-type (WT) cells and 4,458 cells from day 4 post-irradiation samples. Data normalization, dimensionality reduction (PCA and UMAP), clustering, and visualization of cell populations were performed using Seurat.

### Pesudotime and trajectory generation and boxplot signal plotting

#### Pesudotime and trajectory analysis

Pseudotime trajectories were generated using Monocle 2. Single-cell expression data and associated cell-type annotations were imported from the Seurat object into a Monocle CellDataSet. 500 Highly variable genes and fetal and adult Cdx2 associated genes were used as ordering genes, and dimensionality reduction was performed using the discriminative dimensionality reduction with trees (DDRTree) algorithm. Cells were ordered along the resulting trajectory, with the stem-cell population designated as the root, thereby orienting pseudotime from stem cells through differentiated enterocytes. Cell states and differentiation territories were transferred from Seurat object. First component of DDRTree was used to compare the position of WT and radiated samples along the differentiation trajectory.

---

**Figure S1.**
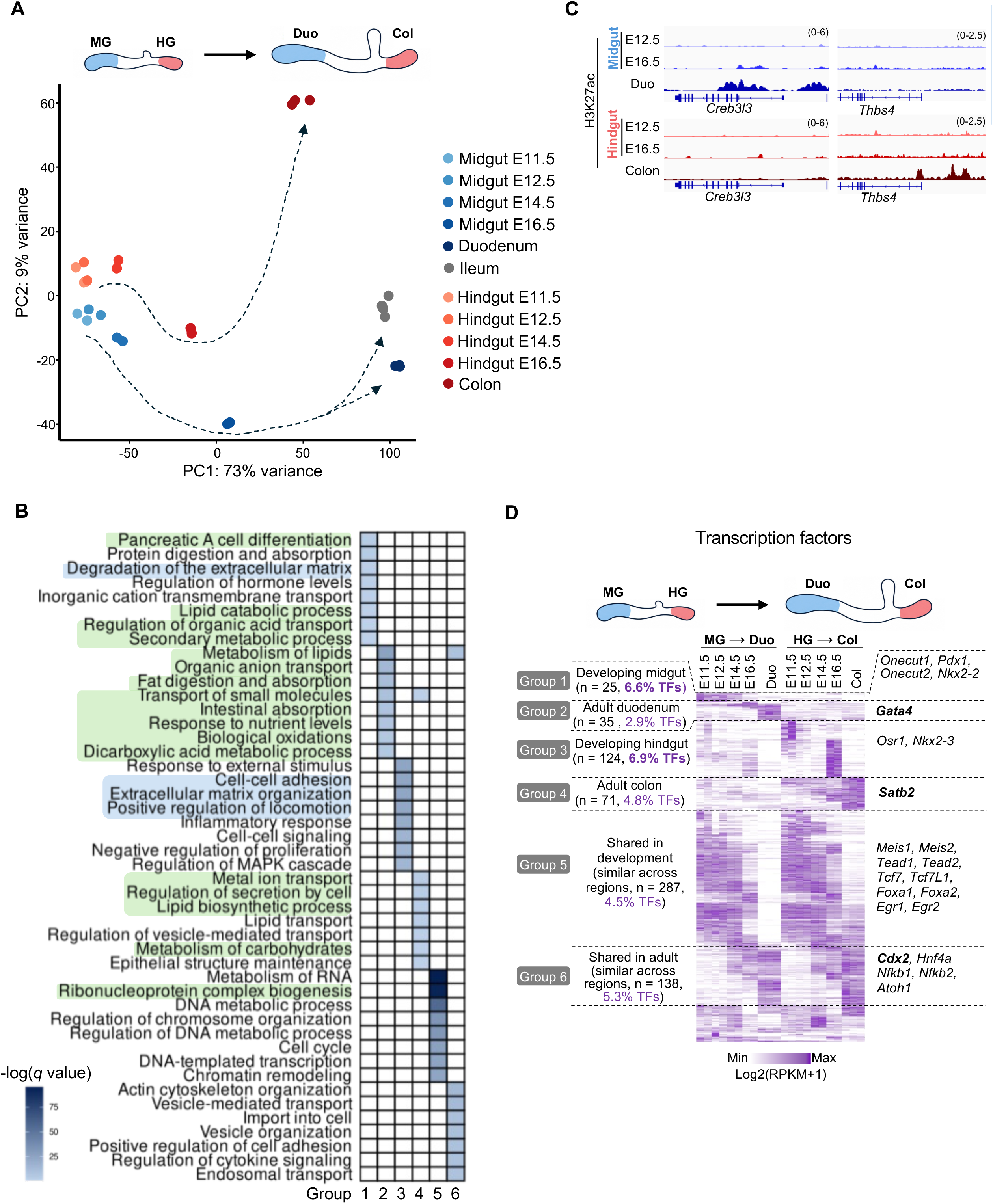
Diversity of regional gene expression programs underlies intestinal structural and functional specialization. (A) Principal-component analysis of gene-expression profiles across embryonic midgut and hindgut development and adult intestinal regions. Samples segregate by regional identity and developmental stage, illustrating progressive divergence of MG- and HG-derived intestinal tissues. Dashed arrows indicate the inferred developmental trajectories from fetal to adult tissues. (B) Gene Ontology biological-process enrichment analysis of genes in Groups 1-6 from Figure 1A. Heatmap intensity represents enrichment significance. Processes associated with epithelial structural development and metabolic function are highlighted in light blue and light green, respectively. (C) Representative genomic locus illustrating H3K27ac signal genes with enhancers at duodenum-specific genes that are absent at the corresponding colon loci, and vice versa. (D) Heatmap of transcription-factor expression dynamics across the MG→duodenum and HG→colon developmental trajectories, within the six gene-expression groups defined in Figure 1A (gene number and TF percentage indicated per group). Representative transcription factors for each group are listed alongside the heatmap; color scale indicates Log2(RPKM+1) expression.

**Figure S2.**
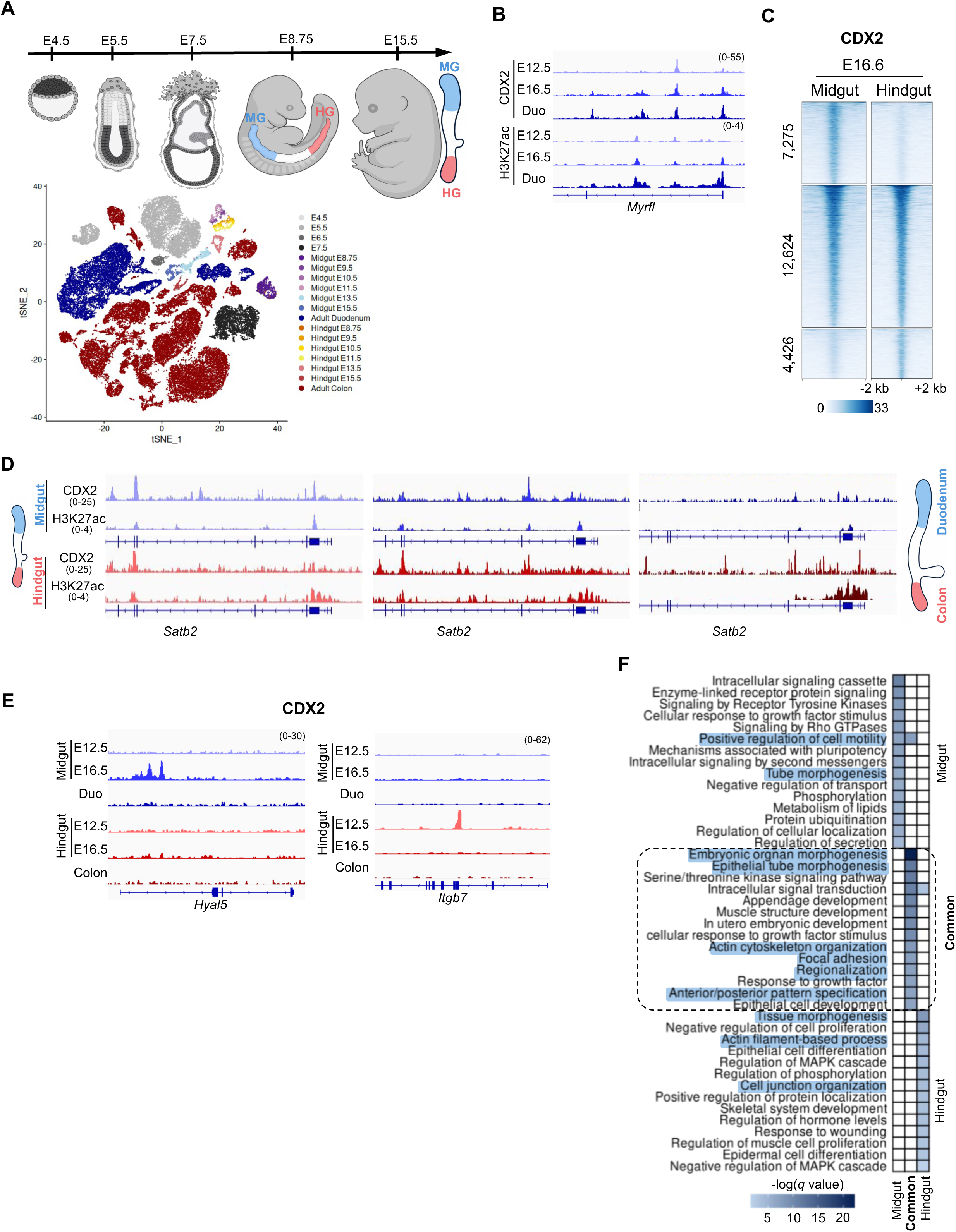
Evolving CDX2 activity underlies development of regional intestinal identity. (A) Schematic of embryonic^42^ and fetal endodermal development^43^, including MG and HG intestinal regions, and adult epithelial cell populations^44^ used to generate the single-cell RNA-seq dataset analyzed in Figure 2A. (B) Representative genome-browser tracks at a midgut gene locus, showing CDX2 and H3K27ac signal at E12.5, E16.5, and in adult duodenum, illustrating the overlap between CDX2 binding and H3K27ac+ enhancer activity. (C) Heatmaps showing CDX2 occupancy at fetal-specific loci that are midgut-specific, hindgut-specific, or shared between both epithelial regions. (D) Representative genomic locus illustrating H3K27ac and CDX2 signal at *Satb2* loci and the transition from shared fetal CDX2 occupancy to region-specific adult binding. (E) Representative genome-browser tracks illustrating CDX2 signal at two distinct CDX2 binding patterns during development: fetal, MG- or HG-specific CDX2 binding that is lost in adult tissue (top), and new, region- and adult-specific CDX2 binding gained in adult tissue (bottom). (F) Gene Ontology biological-process enrichment analysis of genes within ±25 kb of fetal CDX2-bound loci that are common to both midgut and hindgut, or MG-specific or HG-specific, as defined in Figure 2E. Processes related to epithelial cell migration, tissue structure, and morphogenesis (highlighted) are most strongly enriched among genes at commonly bound loci, though also enriched among region-specific gene sets.

**Figure S3.**
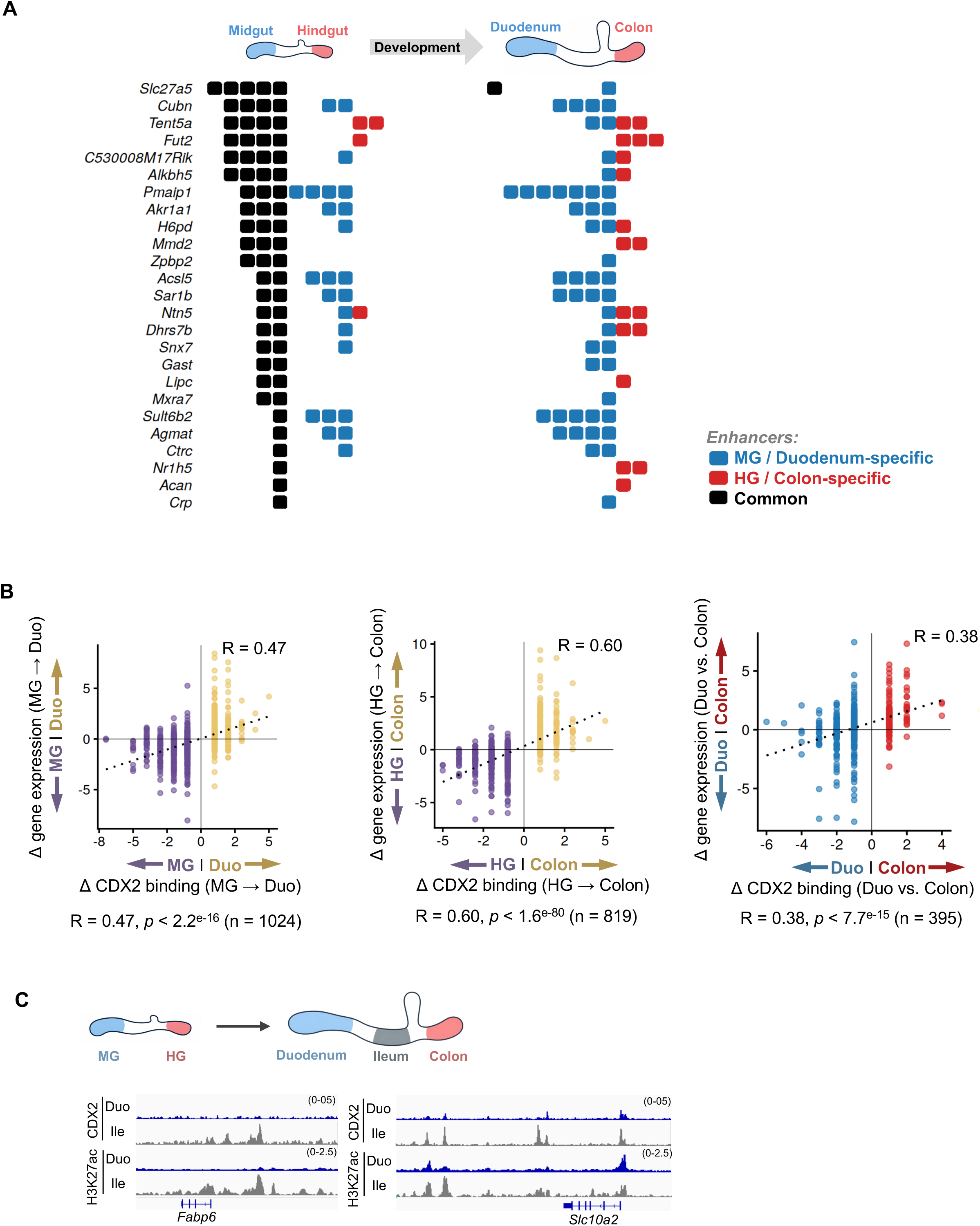
CDX2 binding patterns influence gene expression and co-recruitment of other transcription factors. (A) CDX2 multienhancer occupancy map across fetal MG and HG and their adult duodenal and colonic derivatives. Fetal epithelial states are characterized by extensive multienhancer CDX2 architecture that resolves into adult tissue-specific enhancer configurations associated with regional gene-expression patterns. (B) Correlation between the number of CDX2-bound regulatory elements and expression of linked genes across the indicated fetal, developmental, and adult intestinal regions. (C) Representative genome-browser tracks showing H3K27ac and CDX2 signal at region-specific CDX2 occupancy and enhancer activity, as measured by H3K27ac, in adult duodenum and ileum.

**Figure S4.**
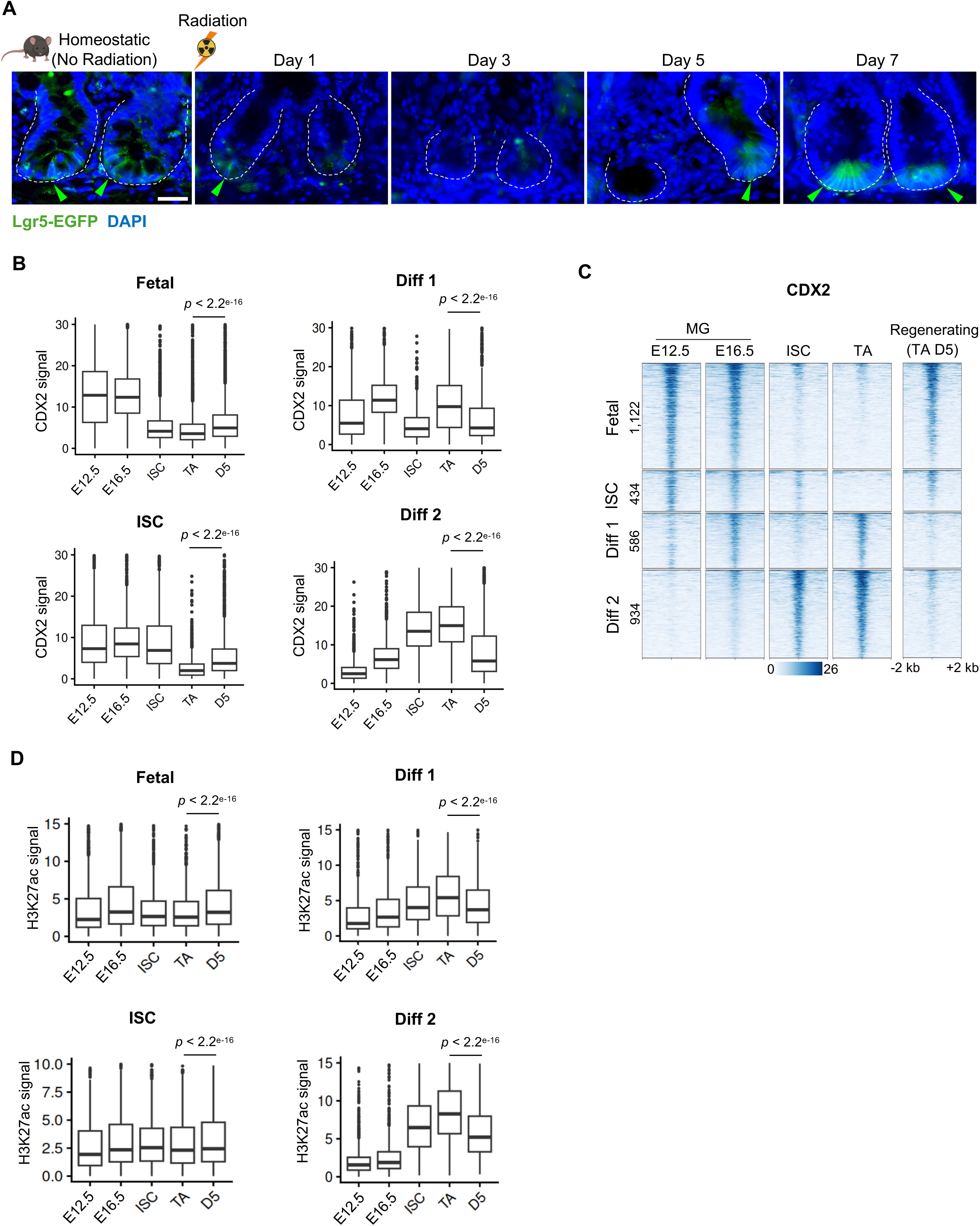
Dynamics of ISC loss and regeneration and associated changes in CDX2 occupancy. (A) Representative fluorescence micrographs showing loss of Lgr5-EGFP+ ISCs following radiation, with maximal depletion at day 3 and recovery by day 5. White dotted lines outline crypts, and green arrowheads indicate crypts containing Lgr5-EGFP signal. Scale bar, 25 μm. (B) Boxplots showing CDX2 occupancy at fetal-, ISC-, and differentiation-associated loci (Diff 1 and Diff 2; Figure 4A) across fetal intestinal development (E12.5 and E16.5), adult epithelial populations (ISC and TA), and regenerating D5 TA cells. Statistical significance was determined using the Wilcoxon signed-rank test (*p* < 2.2 × 10^-16^). (C) Heatmaps showing loci with at least 1.5-fold gain in CDX2 occupancy at fetal- and ISC-associated regulatory elements or at least 1.5-fold loss at Diff 1 and Diff 2 loci in regenerating D5 TA cells compared with homeostatic TA cells. (D) Boxplots showing normalized H3K27ac signal at fetal-, ISC-, and differentiation-associated loci (Diff 1 and Diff 2; Figure 4A) across fetal intestinal development (E12.5 and E16.5), adult epithelial populations (ISC and TA), and regenerating D5 TA cells. Statistical significance was determined using the Wilcoxon signed-rank test (*p* < 2.2 × 10^-16^).

**Figure S5:**
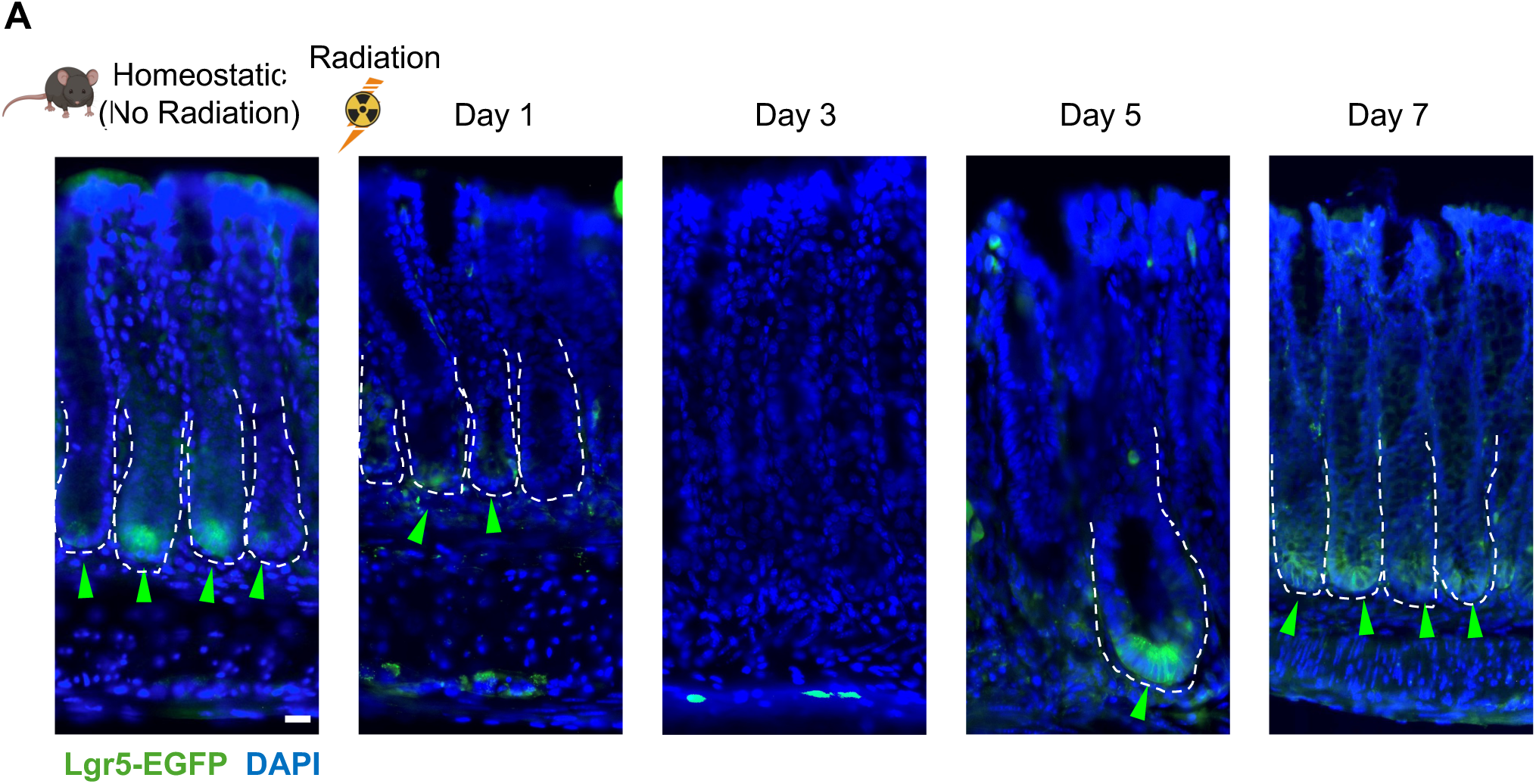
Colonic epithelial regeneration post radiation. (A) Representative fluorescence micrographs showing the sequence of Lgr5+ISC loss post-radiation (day 3), before recovery at day 5 in mouse colon; white dotted lines indicate crypts, green arrowheads indicate crypts with Lgr5-EGFP signal. Scale bar = 25 μm.

**Figure S6:**
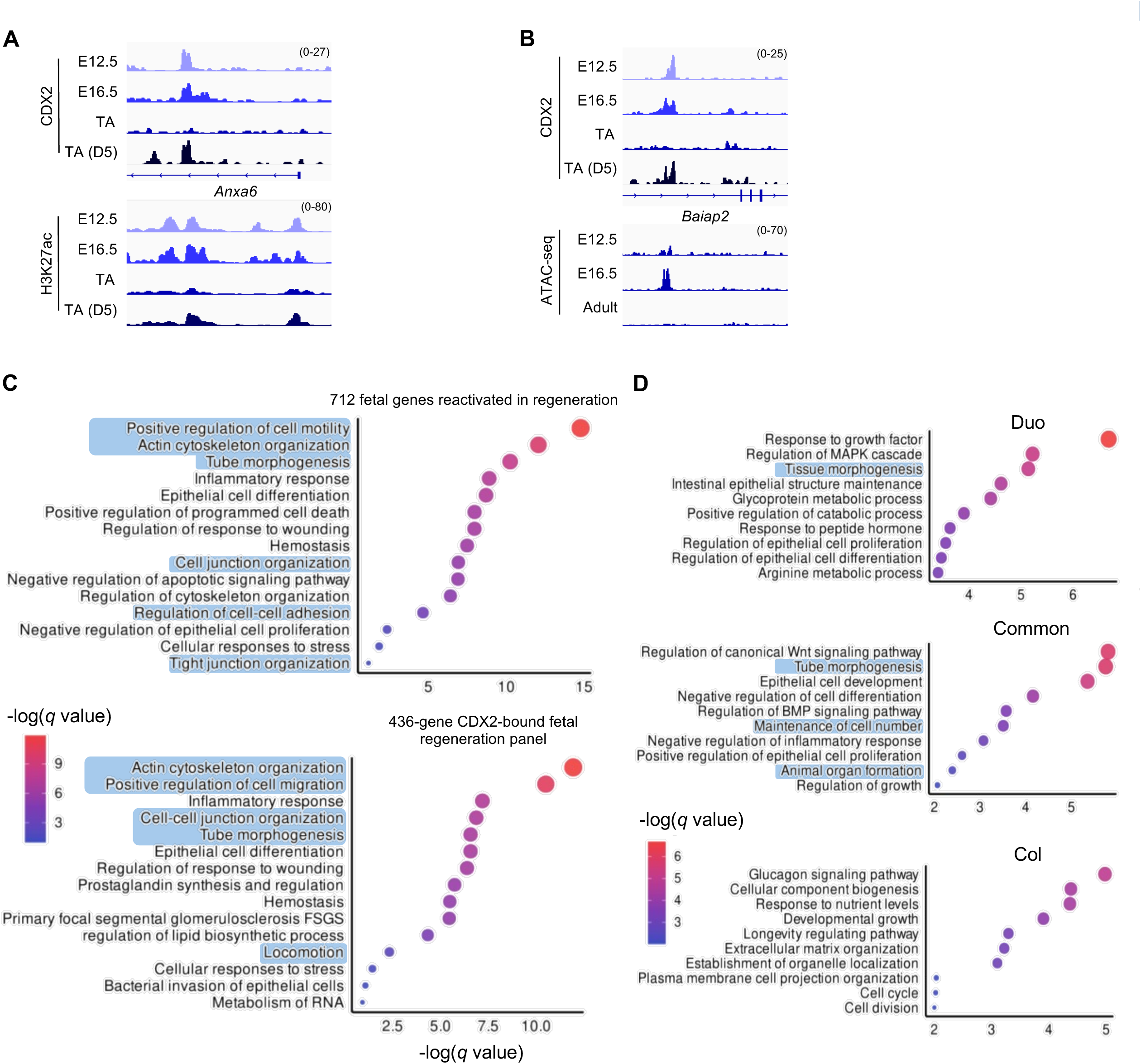
CDX2 based control of epithelial structural patterning in fetal and regenerating tissue. (A) Representative genome-browser tracks of H3K27ac and CDX2 signal illustrating CDX2 binding dynamics at fetal enhancer loci at example published fetal-like genes demonstrating selective rebinding of CDX2 to fetal regulatory elements during epithelial regeneration. (B) Representative genome-browser tracks of ATAC and CDX2 signal illustrating that CDX2 occupancy during regeneration (adult TA D5) is more correlated with chromatin accessibility (ATAC-seq) in fetal (E16.5) cells rather than adult homeostatic cells (TA). (C) Dot plots showing gene ontology analysis of the 712 fetal genes activated across various inductions and injuries, and the 436 CDX2-bound genes among them, reveal enrichment for cell migration and tissue structure/morphogenesis. (D) Dot plots showing gene ontology analysis of genes rebinding CDX2 in a duodenum-specific, colon-specific, or common manner, revealing that commonly bound genes are enriched for cell migration and tissue structure/morphogenesis.

**Figure S7:**
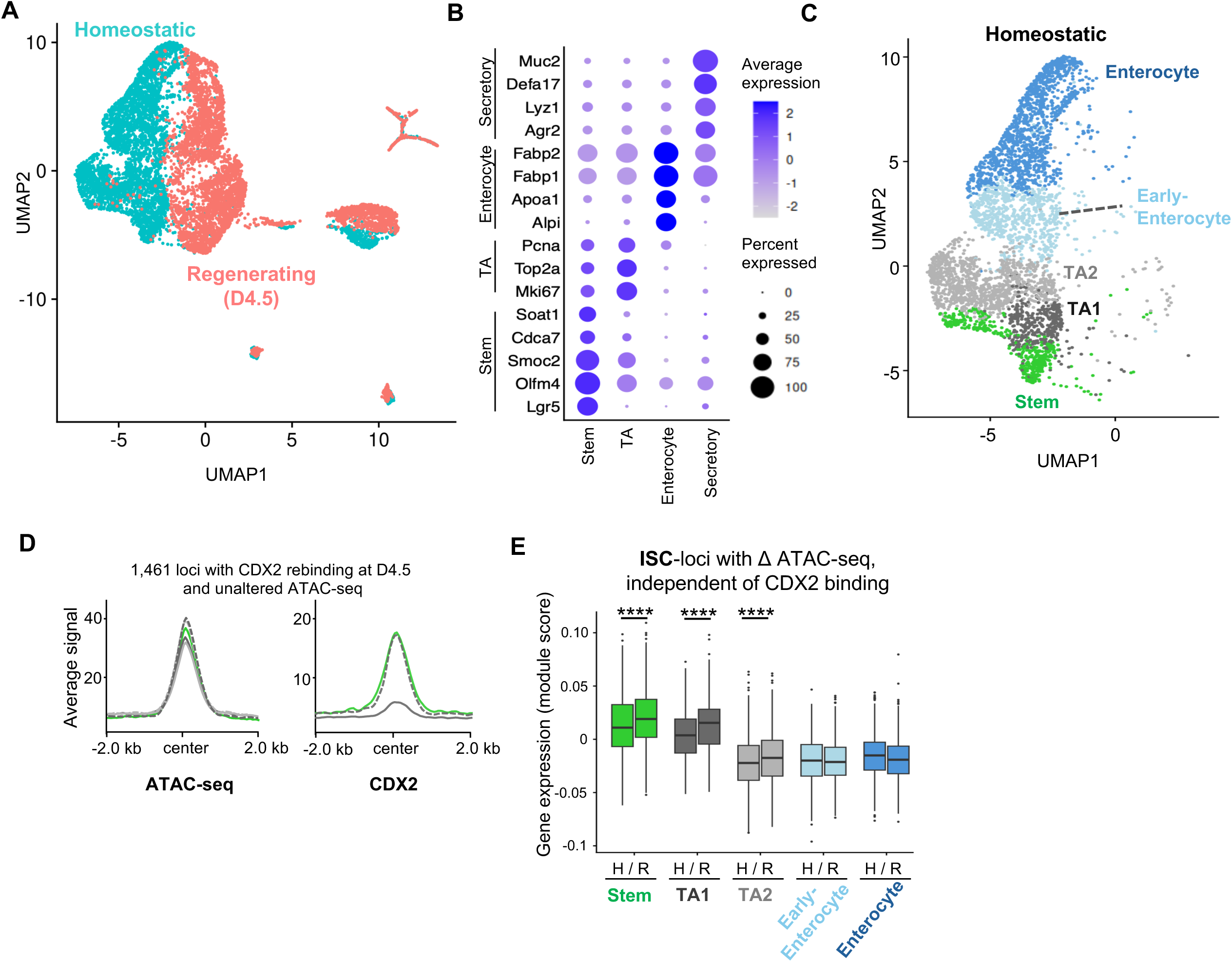
Single-cell Multiome and CDX2 binding analysis together reveal molecular basis of cell state dynamics within regenerating crypts. (**A**) UMAP of scMultiome (scRNA-seq and scATAC-seq) from intestinal epithelium showing homeostatic (cyan) and regenerating (pink) cell populations. UMAP on the right shows all cell states along the ISC → Enterocyte differentiation path in homeostatic (uninjured) and regenerating epithelium (day 4.5 post-radiation); the inset (top-right) identifies homeostatic (cyan) and regenerating (pink) cell populations. (B) Dot plots showing expression levels and percent of cells expressing lineage and cell type marker genes used to define clusters and cell states in the single-cell multiome data. (C) UMAP showing cell states along the ISC → Enterocyte differentiation path in homeostatic (uninjured) epithelium; TA1 cell clusters show lower cycling and higher proximity to ISCs as compared to TA2, and early enterocytes show lower levels of mature enterocyte markers and higher proximity to TA2 populations. (D) Average ATAC-seq (left) and CDX2 (right) signal profiles centered on 1,461 loci with CDX2 rebinding at D4.5 and unaltered ATAC-seq signal. Signal is shown in homeostatic ISC, TA1, and TA2 populations, and in the corresponding regenerating TA1 (D4.5) and TA2 (D4.5) populations. (E) Boxplots showing gene expression scores for genes linked to the 1,355 ISC loci with ATAC-seq signal changes independent of CDX2 binding, in homeostatic (H) and regenerating (R) cells across the five cell states spanning the differentiation trajectory. Center lines indicate medians, boxes indicate interquartile ranges (IQRs), whiskers extend to 1.5 × IQR, and points indicate outliers. Statistical significance was determined using the Wilcoxon rank-sum test (****, *p* < 0.0001).

